# Mitochondrial metabolism is a key determinant of chemotherapy sensitivity in Colorectal Cancer

**DOI:** 10.1101/2024.09.12.611189

**Authors:** Deborah Y. Moss, Connor Brown, Andrew Shaw, Christopher McCann, Nikita Lewis, Aaron Phillips, Sarah Gallagher, William J. McDaid, Andrew Roe, Aisling Y. Coughlan, Brenton Cavanagh, Callum Ormsby, Fiammetta Falcone, Rachel McCole, Scott Monteith, Emily Rogan, Matilda Downs, Sudhir B. Malla, Alexandra J Emerson, Letitia Mohammed-Smith, Shaun Sharkey, Peter F. Gallagher, Arindam Banerjee, Sufyan Pandor, Brett Greer, Christopher Elliott, Aideen Ryan, Philip D. Dunne, Vicky Coyle, Ian G. Mills, Simon S. McDade, Owen Sansom, Triona Ni Chonghaile, Daniel B. Longley, Melissa J. LaBonte, Emma M. Kerr

## Abstract

Therapy resistance is attributed to over 80% of cancer deaths per year emphasizing the urgent need to overcome this challenge for improved patient outcomes. Despite its widespread use in colorectal cancer (CRC) treatment, resistance to 5-fluorouracil (5FU) remains poorly understood. Here, we investigate the transcriptional responses of CRC cells to 5FU treatment, revealing significant metabolic reprogramming towards heightened mitochondrial activity. Utilizing CRC models, we demonstrate sustained enhancement of mitochondrial biogenesis and function following 5FU treatment, leading to resistance in both in vitro and in vivo settings. Furthermore, we show that targeting mitochondrial metabolism, specifically by inhibiting Complex I (CI), sensitizes CRC cells to 5FU, resulting in delayed tumour growth and prolonged survival in preclinical models. Additionally, our analysis of patient data suggests that oxidative metabolism signatures may predict responses to 5FU-based chemotherapy. These findings shed light on mechanisms underlying 5FU resistance and propose a rational strategy for combination therapy in CRC, emphasizing the potential clinical benefit of targeting mitochondrial metabolism to overcome resistance and enhance patient outcomes.

## Main

Therapy resistance is a major clinical problem, limiting effectiveness of cancer treatments. Resistance is promoted by a range of factors, present in the tumour prior to therapy (described as intrinsic resistance) or selected for during therapeutic exposure (termed acquired resistance). These factors can stem from the tumour cells themselves, the surrounding tumour microenvironment or systemic influences^1–3^. Understanding the molecular determinants of resistance and using this to guide rational combination therapy approaches is fundamental to improve clinical outcomes for cancer patients.

Since its discovery in the 1950s, the anti-metabolite 5-fluorouracil (5FU) has been the cornerstone of fluoropyrimidine (FP)-based chemotherapy treatment for colorectal cancer (CRC) patients. Given as a single agent, it promotes modest increases in survival, and these effects are enhanced when combined with Oxaliplatin and/or Irinotecan (FOLFOX, FOLFIRI)^4–7^. However, despite being the mainstay of CRC chemotherapy treatment, the primary anti-tumour mechanisms, and more importantly, mechanisms facilitating 5FU resistance and persistence remain poorly defined. 5FU’s multi-faceted mechanism of action, causing direct damage to DNA (FdUTP) and RNA (FUTP) as well as the DNA and metabolic impacts caused by inhibition of thymidylate synthase (TS) via FdUMP^8,9^, makes it more difficult to identify rational combination partners that would enhance clinical responses without eliciting adverse systemic toxicity.

Reprogramming of cancer cell energetics is fundamental to tumour progression and spread^10,11^, but is also critically linked to therapeutic efficacy^12–15^. Targeting such metabolic programs presents novel opportunities to enhance drug responses and improve patient outcomes^12,16–18^. Therefore, a better understanding of the metabolic programs critical in supporting cancer cell survival following 5FU treatment is required to rationally design more effective FP-based chemotherapy combination regimens.

## Results

Using the well characterised CRC cell model, HCT116, we profiled dynamic transcriptional responses to 5FU treatment prior to the onset of significant growth arrest and apoptosis (24-48 hr). Differential gene expression analysis using the DESeq2 package identified 1856 and 3350 differentially expressed genes (DEG, adj. p<0.05) at 24 hr and 48 hr, respectively (Ext. data figure 1a). Gene Set Enrichment Analysis (GSEA) shows differential signalling programs in a number of MSigDB Hallmark biological networks, including several metabolic programs (Ext. data figure 1b). When DEG’s were cross-referenced to a metabolic gene list (2921 genes) compiled from previously published studies^10,19^, we observed significant over-representation of metabolic DEG, as determined by Fisher’s exact test (Ext. data figure 1a). Interestingly, the pattern of change in these metabolic genes (280 and 550 at 24 hr and 48 hr, respectively, (Ext. data figure 1c) suggest that the metabolic response is mounted quickly, but sustained and enhanced over time, given that ∼75% of the metabolic DEG at 24 hrs are also found at 48 hrs (Ext. data figure 1c). Further examining the change in gene expression revealed an extensive number of these metabolic genes are increased in expression both at 24 (Ext. data figure 1d) and 48 hrs (Ext. data figure 1e). Indeed, the directionality of change (i.e. increased or decreased expression) indicates that expression genes associated with metabolism, are predominantly increased, and thus we postulate metabolic activity may be enhanced in response to 5FU treatment (Ext. data figure 1f). Furthermore analysis of curated metabolic gene sets (Ext. data figure 1g) suggests significant changes in expression of gene-sets related to a range of metabolic signalling networks, including lipid, protein, nucleotide, carbon and Reactive Oxygen Species (ROS) metabolism (Ext. data figure 1g). Together, these transcriptional analyses suggest that 5FU results in a strong and sustained impact of expression of metabolism related genes that has potential to significantly impact in cancer cells and warranted further exploration experimentally. Therefore, we next sought to assess 5FU-driven metabolic reprogramming in CRC models.

### 5FU treatment alters mitochondrial biogenesis and dynamics to increase activity

The mitochondria are a unique organelle, coordinating metabolic and apoptotic crosstalk to tightly regulate cell survival^20,21^. Given the breadth of metabolic impact we uncovered by RNASeq analysis (Ext. data figure 1), we examined the impact of 5FU treatment on this metabolic hub. In cells that survive treatment, 5FU promotes a sustained increase in mitochondrial content over 72 hrs as shown by changes in levels of NonylAcridine Orange (NAO) staining (Figure 1a), mitochondrial gDNA (relative to total gDNA/cell, Figure 1b) and TOM20 protein expression (Figure 1c) when compared to PBS control (CTRL). This was a temporal response to 5FU treatment with sustained increases observed over time (Ext. data figure 2a-b). These mitochondria were functional, as reflected by enhanced Electron Transport Chain (ETC) protein expression (Figure 1c) and activity (Figure 1d and Ext. data figure 2c). Increased ETC activity was further supported by enriched mitochondrial metabolism, illustrated by an increase in ^13^C_6_-glucose incorporation into TCA cycle metabolites and glutathione production following 5FU treatment (Figure 1e). Importantly, these changes in mitochondrial mass are not related to engagement of apoptosis, since inhibition of caspase activity using zVAD-fmk did not prevent increases in mitochondrial abundance per cell (Ext. data figure 2d). Using electron microscopy, we analysed the effects of 72 hrs 5FU treatment on mitochondrial morphology to identify potential organelle dysfunction. Upon examination, both the 5FU-treated and control groups exhibited comparable mitochondrial features. There was no noticeable swelling or rupture, and the appearance and abundance of cristae were consistent between the two groups (as illustrated in Figure 1f and Ext. data figure 2e). However, looking at the aspect ratio (defined as the ratio of the major:minor axes lengths) we observed an elongation of the mitochondria (Figure 1f), which is again indicative of a more active organelle^22,23^. These changes were accompanied by altered expression of the proteins that regulate mitochondrial morphology^24^. We observed a modest increase in OPA1 expression (∼1.25-1.6 fold) and a considerable decrease in DRP1 expression (∼5-10 fold) 72 hrs post 5FU treatment (Figure 1g), which correlated with increased elongation of the mitochondria. We confirmed no change in cytochrome C release at baseline in cells surviving 5FU treatment, or indeed any mitochondrial apoptotic priming based on BH3-profiling^25,26^ induced by 5FU treatment (Ext. data figure 2c). This was corroborated by observation of a modest transient dip in mitochondrial membrane potential at 48 hrs that fully recovered by 72 hrs. Taken together, these data demonstrate that CRC cells capable of enduring 5FU treatment *in vitro* have significant, sustained reprogramming of mitochondrial metabolism that drives enhanced mitochondrial biogenesis and sustained mitochondrial fission to facilitate increased mitochondrial activity, thereby promoting cell survival.

**Figure 1:**
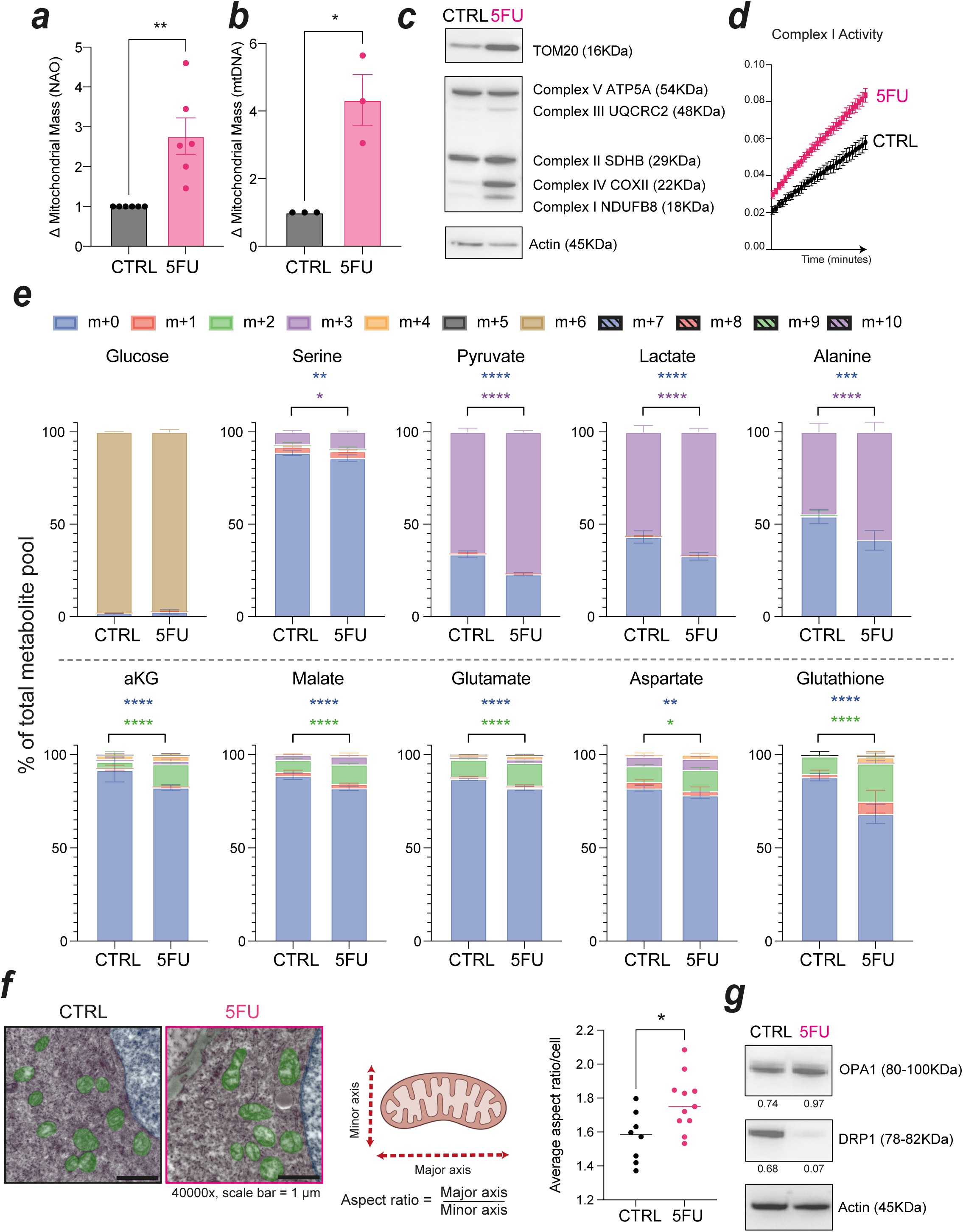
5FU treatment alters mitochondrial phenotypes. Mitochondrial mass in HCT116 cells treated ± 5FU for 72 hrs measured by Nonyl-Acridine Orange (NAO) staining (a) or mitochondrial DNA content (b), data points show biological replicates from triplicate means, normalised to mean negative control and vehicle CTRL from technical replicates. Bars show biological mean, ± s.e.m.; (c) Western blot analysis of HCT116 ± 5FU protein lysates; (d) CI kinetic activity assay in HCT116 cells treated ± 5FU, data points show mean of technical replicates ± s.d., representative data shown of 3 independent experiments; (e) LC-MS metabolomics analysis of labelling in metabolites downstream of ^13^C_6_-glucose from HCT116 cells ± 5FU. Isoform abundance as percentage (%) of total pool is indicated in stacked bar charts, data shows mean of 5 replicates ± s.d., representative of two individual experiments; (f) transmission electron microscopy images of HCT116 cells ± 5FU, with the measurements taken from visible mitochondria to calculate aspect ratio illustrated. Average aspect ratio per cell shown, data points show median of technical replicates. Scale shown = 1 µm ; (g) Western blot analysis of HCT116 ± 5FU protein lysates, with expression relative to Actin denoted below each blot. ****p<0.0001, ***p<0.001, **p<0.01, *p<0.05, calculated by paired T-test (a-b) Two-way anova with Šídák’s multiple comparisons test (e) or unpaired T-test (g).

### Mitochondrial phenotypes are recapitulated in other KRAS mutant CRC models

HCT116 cells are an extensively utilized human colorectal model that carry a *KRAS* G13D mutation but retain wild-type *p53* and *APC*^27,28^. KRAS signalling has been linked to enhanced mitochondrial metabolism by various mechanisms in colorectal, lung and pancreatic cancers^29–32^, as has APC and p53^33–37^. We next sought to confirm our observations on the mitochondrial impact of 5FU in more complex models of CRC. Using *VilCre^ERT2^/Apc^Fx^/Kras^G12D^/p53^Fx^* genetically engineered mouse models (GEMM)^38^ we examined the effects of 5FU treatment *in vitro* using 3D organoid cultures or *in vivo* using multiple methods of tumour induction.

First, organoids were isolated from the colon of *VilCre^ERT2^;Apc^Fx/Fx^;Kras^G12D/+^;p53^Fx/Fx^*(*AKP)* mice 72 hrs post-tamoxifen intraperitoneal (i.p.) injection and cultured as previously reported (Figure 2a)^38,39^. Organoids displayed similar 5FU sensitivity to human models of CRC, with GIC_50_ (50% Growth Inhibitory Concentration) values of 1-10 µM (*data not shown*). *AKP* organoids were treated with 5FU or PBS control (CTRL) for 72 hrs and metabolic gene and protein expression was quantified. Quantitative real-time PCR (qPCR) analysis of a panel of metabolic genes identified in RNA-seq analysis (Ext. data figure 1) demonstrated that the metabolic impact of 5FU treatment observed in our human model was recapitulated in these murine organoids (Figure 2b). Furthermore, we again found that these transcriptional changes are reflected at the protein level with enhanced TOM20, and Complex I (CI) and IV (IV) expression observed (Figure 2c). These data also suggest that the mitochondrial component of the metabolic impact of 5FU is not p53 or Apc mediated. Importantly, it also demonstrates that the phenotypes observed in HCT116 models are not limited to 2D culture^40^.

**Figure 2:**
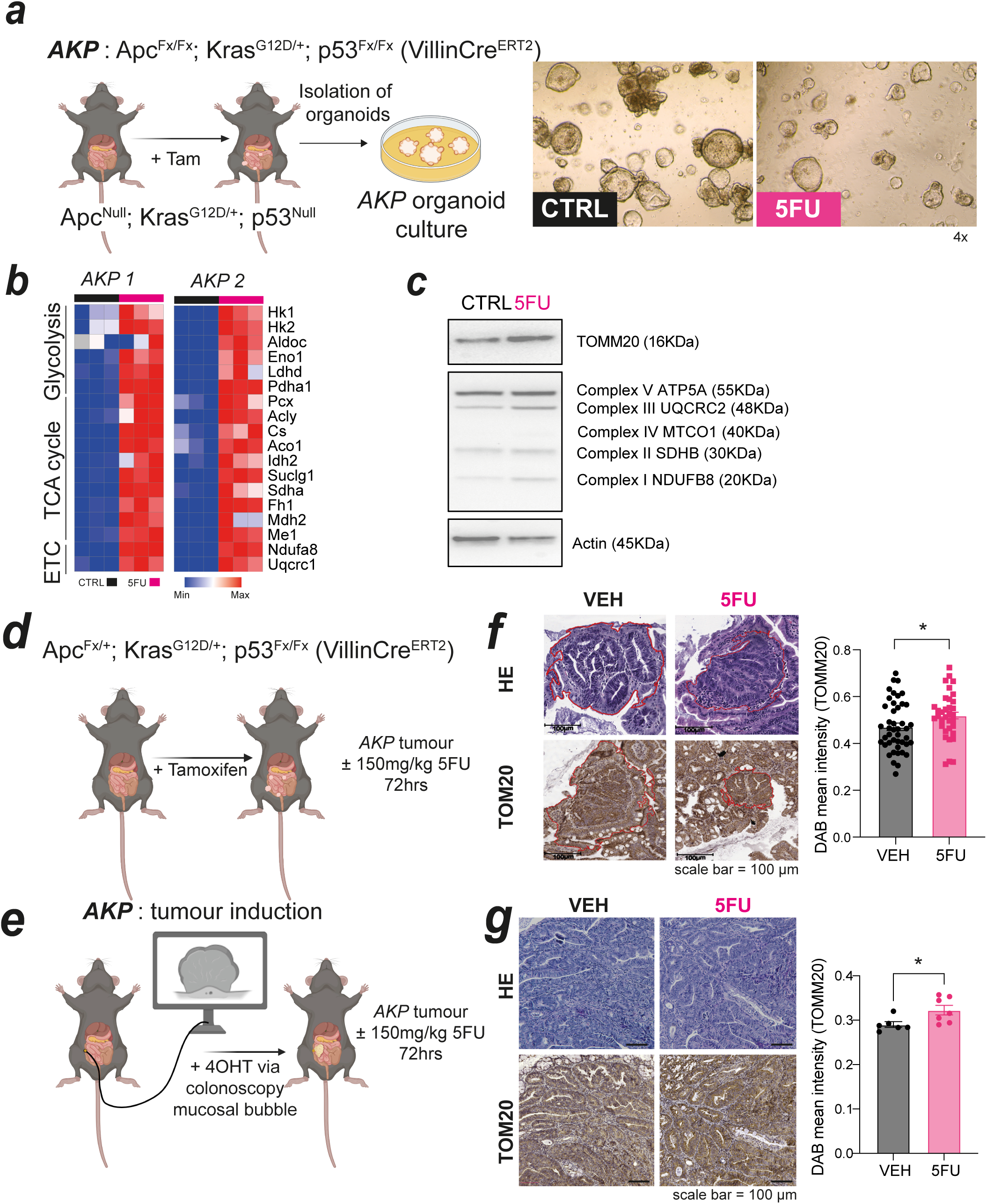
Mitochondrial impact of 5FU is recapitulated in murine models of CRC. (a) Schematic illustrating isolation of *Apc^Null^;Kras^G12D/+^;p53^Null^* (*AKP*) organoid cultures and sensitivity to 5 μM 5FU treatment after 72 hrs. Images taken at 4x magnification; (b) Heatmap showing change in metabolic gene expression measured by qPCR analysis of mRNA from two independent *AKP* organoid lines ± 5FU; (c) Western blot analysis of protein lysates from *AKP* organoids ± 5FU; Schematic illustrating 5FU experimental treatments in Apc^Fx/+^;Kras^G12D/+^;p53^Fx/Fx^ (VillinCreERT2) mice (d) or *AKP* mice (e); (f-g) Representative H&E and TOM20 images in mice from (d) and (e) ± 5FU treatment, with quantification of TOM20 expression per tumour by QuPath analysis shown ± s.e.m. Scale shown = 100 μm. *p<0.05 by unpaired t-test.

We next sought to examine this mitochondrial biogenesis phenotype in whole tumour ecosystems to ensure our observations were not linked to a culture phenomenon as has previously been described for other metabolic reprogramming studies^41^. We used i.p. injection of tamoxifen (Figure 2d, *Apc^Fx/+^* mice) or orthotopic endoscopy-guided intramucosal injection^42–44^ of 4-OHT (Figure 2e, *Apc^Fx/Fx^* mice) to drive allelic recombination and promote tumour formation. Tumours were allowed to develop for 3-6 weeks until mice showed signs of tumour burden, at which point mice were randomised and treated with a single i.p. injection of 150 mg/kg 5FU or PBS vehicle for 72 hrs before mice were culled and tumour tissue was collected (Figure 2d-g). In both low-grade lesions (Figure 2f) and high-grade tumours (Figure 2g), we observed increases in mitochondrial content following 5FU treatment as was observed *in vitro,* demonstrated by the increase in TOM20 staining (Figure 2f-g). These *in vitro* and *in vivo* data demonstrate that 5FU treatment robustly increases mitochondrial content in colorectal tumour cells.

### TS inhibition by 5FU regulates mitochondrial impact

5FU has a multimodal mechanism of action: promoting DNA damage directly via its metabolite FdUTP or by inhibition of TS via FdUMP; by depleting the pool of thymidine available for DNA synthesis; and RNA damage via FUTP (Figure 3a)^8^. 5FU has previously been demonstrated to have a significant metabolic impact shortly after treatment, rapidly and consistently inhibiting TS^9^. Indeed, in our setting we can recapitulate this inhibition of TS and further confirm that this inhibition persists beyond the previously reported 6 hr timepoint through the continued accumulation of dUMP at 24 hrs (Figure 3b), correlating with our observed persistent metabolic alterations at the transcriptional (Ext. data figure 1) and the mitochondrial level (Figure 1, Ext. data figure 2). Given this sustained increase in dUMP, we tested if it was inhibition of TS in cells surviving 5FU treatment that promoted the observed reprogramming of mitochondria towards increased biogenesis and activity. We tested this using genetic and pharmacological inhibition of TS activity. Using siRNA against *TYMS* we show that knockdown of TS increases mitochondrial content in cells, indicated by NAO staining (Figure 3c) and TOM20 expression (Ext. data figure 3a). We validated these data using the antifolate specific TS inhibitor, Raltitrexed^45^. In line with our hypothesis, Raltitrexed also significantly increased mitochondrial content (Figure 3d), as can treatment with other TS-targeted antifolates, Pemetrexed and Methotrexate in colon (Ext. data figure 3b) and lung cancer cell lines (Ext. data figure 3c). Together, these data demonstrate that the observed changes in mitochondrial abundance caused by 5FU are mediated through it’s inhibitory action on TS.

**Figure 3:**
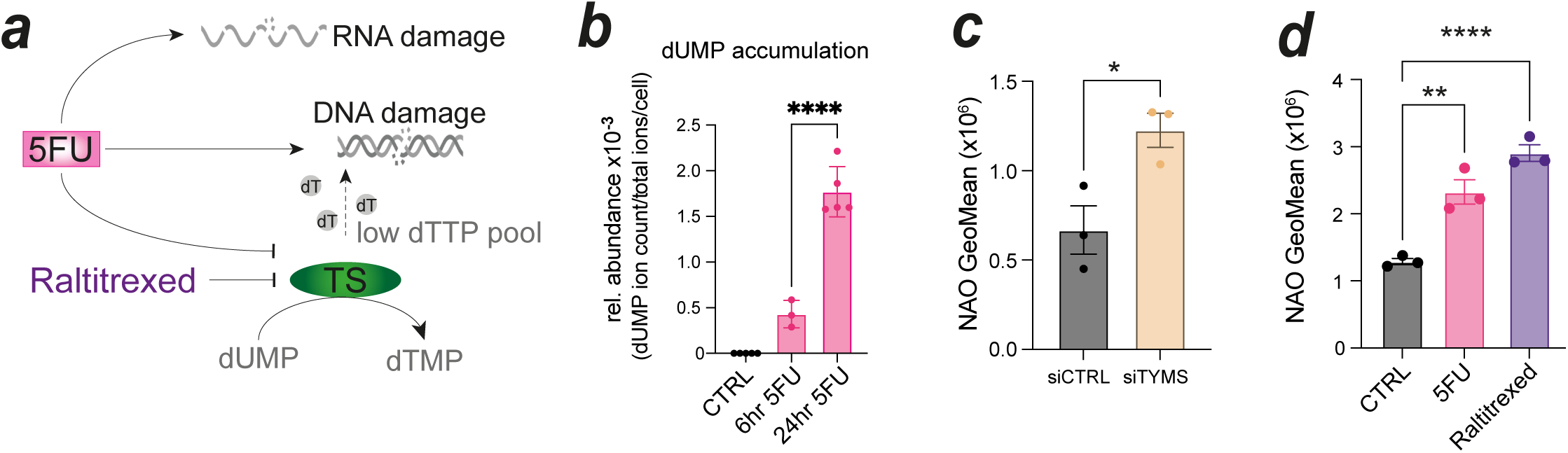
Thymidylate Synthase inhibition drives increased mitochondrial mass. (a) Schematic showing impact of 5FU and Raltitrexed treatment; (b) relative intracellular dUMP abundance normalised to total ion count/cell in HCT116 cells treated with 5FU for 6 and 24 hrs. Representative data shows technical replicates ± s.d. (n=2); (c) Mitochondrial mass assessed by NAO staining in HCT116 cells transfected with siRNA against CTRL or TYMS (c) or treated with 5FU or Raltitrexed (d) for 72 hrs. Data shows biological replicates ± s.e.m.; ****p<0.0001, ***p<0.001, **p<0.01, *p<0.05, by one-way ANOVA with Tukey’s (b) and Dunnett’s (d) multiple comparisons test or t-test (c).

### Mitochondrial metabolism facilitates 5FU resistance in CRC models in 2D and 3D

Clinically, CRC patients usually undergo 6-12 cycles of 5FU- or fluoropyrimidine (FP)- based chemotherapy^46,47^. However, after initial sensitivity, many patients develop resistance to the treatment^48,49^. We postulated that a change in mitochondrial metabolism could relate to 5FU sensitivity and/or resistance. To demonstrate cause and effect, we determined whether modulating mitochondrial metabolism was sufficient to promote 5FU resistance. To test this, we cultured HCT116 cells in DMEM media where glucose (25 mM) was replaced with galactose (25 mM), a method previously described to promote enhanced TCA cycle flux (Figure 4a)^50,51^. When compared with glucose-control cells, cells cultured in galactose were ∼2.5 times less sensitive to 5FU (Figure 4b).

**Figure 4:**
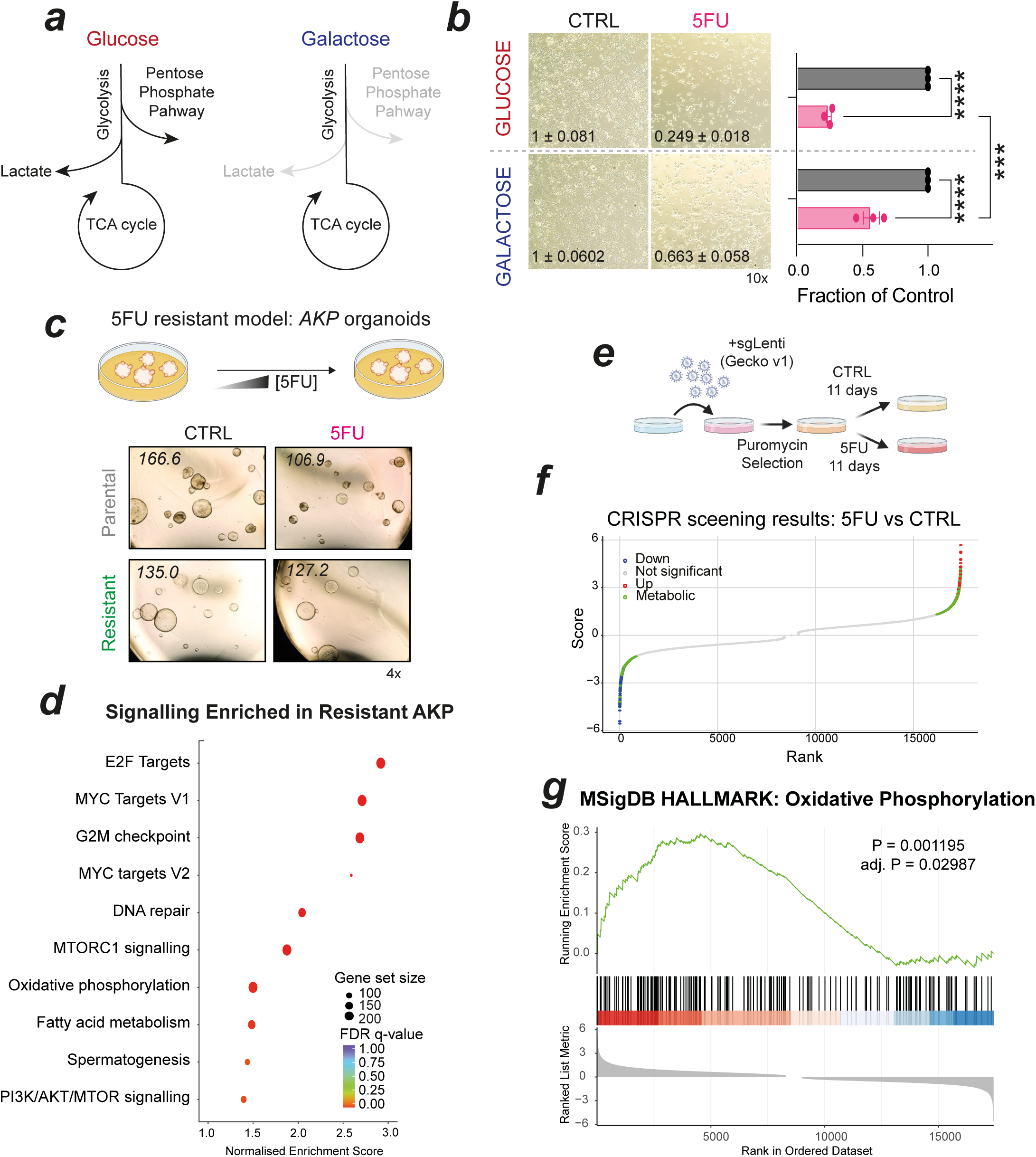
Mitochondrial metabolism dictates sensitivity to 5FU. (a) Schematic representing selectivity of Galactose culture to enhance TCA cycle; (b) Representative images of HCT116 cells cultured in DMEM + 25mM Glucose or Galactose as indicated ± 5FU (magnification 10x). Quantification of n=3 experiments, data points indicate biological replicates ± s.e.m. normalised to media control (CTRL); (c) Schematic and representative images illustrating generation of 5FU resistant *AKP* organoids (magnification 4x), with mean organoid diameter denoted; (d) GSEA analysis of RNASeq data from Parental and 5FU Resistant (Resistant) *AKP* organoids; (e) Schematic representing experimental design of CRISPR screen in HCT116 cells; (f) dot plots showing gene Rank against Score of 5FU vs control (CTRL) treated cells, with significant genes (±1.5xSD) denoted, and metabolic genes highlighted; (g) GSEA analysis of MSigDB Hallmark Oxidative Phosphorylation signature using data from (f) as input, with P and adj. P value reported.***p<0.001, ****p<0.0001 by two-way ANOVA with Šídák’s multiple comparisons test.

We next examined if promoting 5FU resistance modulated mitochondrial metabolism. We generated *AKP* organoids that were ∼4x more resistant to 5FU treatment using chronic exposure to increasing 5FU doses over 6-9 months. When compared to passage matched parental AKP models which retained sensitivity to 5FU (Figure 4c), resistant organoids continued to grow in the presence of 5FU treatment. Transcriptional profiling of these resistant organoids show increases in oxidative phosphorylation and fatty acid metabolism-related genes relative to parental lines (Figure 4d) demonstrating enhanced mitochondrial signalling in the context of acquired 5FU resistance. We confirmed these observations in an existing HCT116 model of acquired 5FU resistance first published in 2004^52^. Again, generated using continuous culture in increasing concentrations of 5FU (Ext. data figure 4a), these HCT116 resistant cells (HCT1165FUR) exhibit an ∼4.5-fold reduction in sensitivity to 5FU. HCT1165FUR cells display increased ETC gene expression (Ext. data figure 4b, qPCR) and mitochondrial metabolite abundance (Ext. data figure 4c, LC-MS). Orthogonally, in a HCT116 5FU-CRISPR screen, where cells infected with the Gecko v1 library were cultured ± 5FU for 11 days (Figure 4e), the analysis showed that ∼7.3% of significant genes were metabolic (Figure 4f), with significant enrichment for OxPhos genes (Figure 4g), providing an independent validation of these observations. Together, these data allow us to conclude that increased mitochondrial metabolism promotes 5FU persistence and resistance, resulting in treatment failure.

### Oxidative metabolism signatures predict response to 5FU-based chemotherapy treatment

Given the observations in the acquired resistance models, we next postulated that elevated mitochondrial metabolism may be a predictor of clinical response to 5FU-based chemotherapy. As 5FU is most often administered in combination with other DNA damaging chemotherapies, including FOLFOX (5FU/Oxaliplatin) or FOLFIRI (5FU/Irinotecan) regimens, we also tested whether the same metabolic/mitochondrial responses were elicited in response to combination treatments. In HCT116 cells treated with FOLFOX (5FU + 1 µM Oxaliplatin) or FOLFIRI (5FU + 5nM SN38, active metabolite of Irinotecan), we observed upregulation of mitochondrial mass (Ext. data figure 5a), mitochondrial activity (Ext. data figure 5b) and ETC complex protein expression (Ext. data figure 5c), suggesting that the mitochondrial impact of combination treatments is driven by 5FU and is maintained in these combination treatments.

Confident that mitochondrial metabolism also plays a role in facilitating survival following FOLFOX and FOLFIRI treatments, we next examined a cohort of CRC patients treated with adjuvant FP-based chemotherapy. This cohort, known as the Taxonomy cohort^53^, is comprised of 156 stage II/III patients, 66 of whom went on to receive adjuvant 5FU-based chemotherapy (illustrated in Figure 5a). Tumour resection (chemo-naïve) samples were previously transcriptionally profiled and data are available from GEO (GSE103479). Transcriptional data from these 66 treated patients was first analysed by GSEA, comparing good outcome patients (top 33% who had best overall survival) with poor outcome patients (bottom 33% who had poorest overall survival). Here, analyses showed that metabolic pathways again ranked highly in poor outcome patients (Figure 5b). Specifically, fatty acid metabolism (NES = 1.66), Oxidative Phosphorylation (NES = 1.60, OxPhos) and mTORC signalling (NES = 1.48) ranked as top three pathways enriched in patients with poorer outcomes to 5FU-based treatment. Given our data with acquired resistance, we next assessed if OxPhos signalling (as an indicator of mitochondrial metabolism) alone was sufficient to predict response to 5FU-based chemotherapy. Using single sample GSEA (ssGSEA), we assigned each patient an OxPhos score using the Hallmarks Oxidative Phosphorylation signature^54,55^ and then split patients based on the cohort mean (Figure 5c). Those above the mean were considered OxPhos High, and those below as OxPhos Low. We compared survival outcomes between the two groups and observed a significant difference in overall survival (Figure 5d, p=0.0387, Hazard ratio=3.036). Importantly, when we compared OxPhos signalling in the same way in patients who did not receive 5FU-based chemotherapy treatment in the same cohort, we observed no difference in survival (Ext. data figure 6), linking differences in patient survival directly to 5FU response. These data again demonstrate that high OxPhos is associated with 5FU resistance and suggest that enhanced mitochondrial metabolism promotes treatment failure clinically.

**Figure 5:**
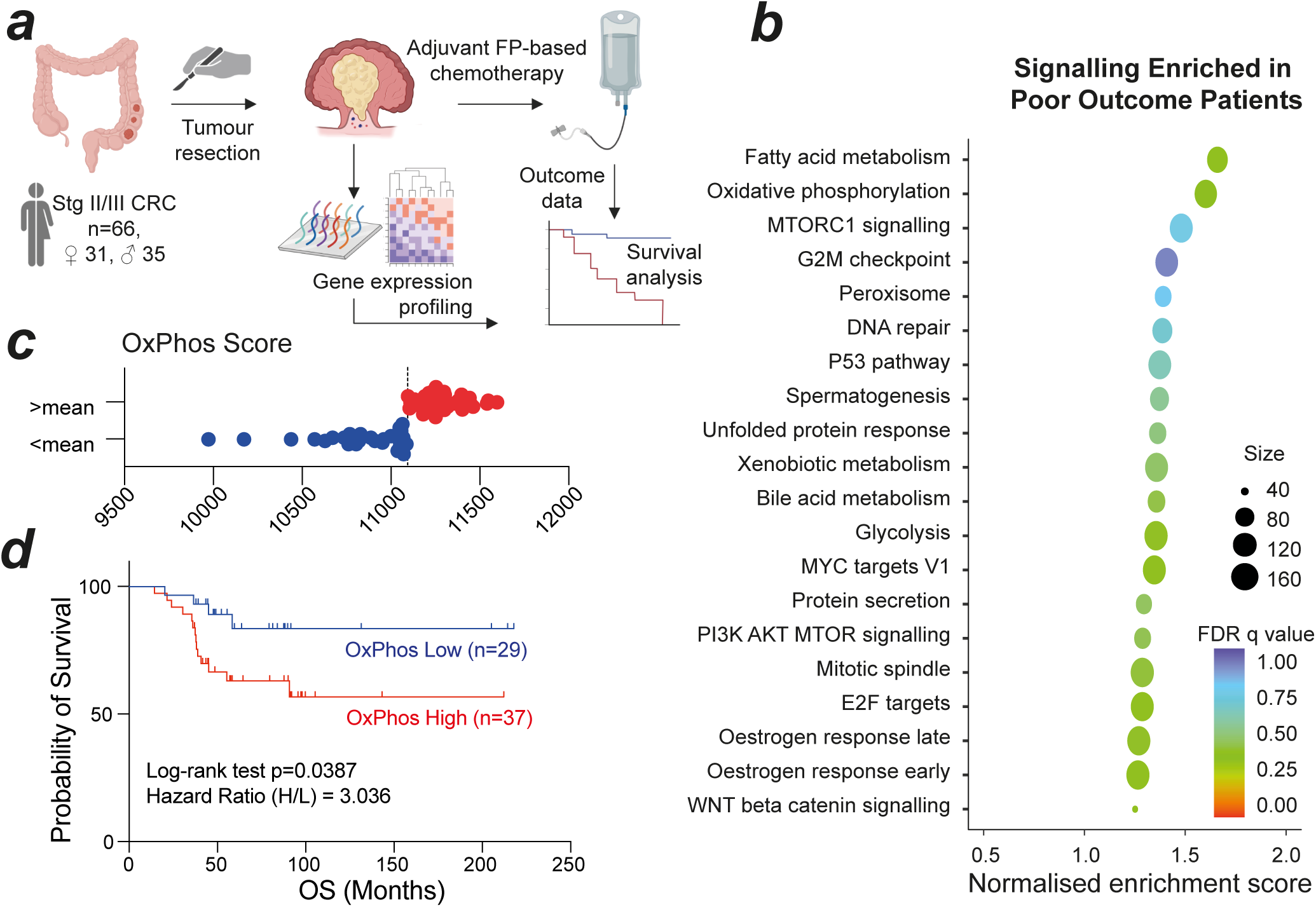
Mitochondrial metabolism predicts sensitivity to adjuvant 5FU-based chemotherapy. (a) Schematic illustrating the Taxonomy cohort samples used for analysis; (b) GSEA analysis of upper vs lower tertiles ranked on overall survival, graph shows Normalised Enrichment Score (NES) for Poor Outcomes patients; (c) OxPhos scores per patient, split by cohort mean; (d) Kaplan Meier Overall Survival (OS) of patients split as in (c), with Log rank test and Hazard Ratio analyses reported.

### Targeting ETC CI enhances 5FU response and extends survival

Up to this point, we have shown that 5FU treatment alters mitochondrial biogenesis and dynamics and that 5FU-resistant cancer cells display an upregulation of mitochondrial metabolism *in vitro* and clinically. The ultimate goal of better understanding 5FU persistence and resistance mechanisms is to allow us to design rational treatment combinations that are more effective than the current standard of care. The increase in mitochondrial mass and activity observed in response to 5FU (Figures 1 & 2) suggests that this adaption to cellular stress may in fact promote an imposed dependency on such mitochondrial signalling that we could therapeutically exploit. Given the increases in ETC CI expression (Figure 1b) and activity (Figure 1c), we postulated that pharmacological inhibition of CI would prove useful in enhancing 5FU-driven anticancer effects. We tested the impact of two clinically relevant inhibitors: IACS-010759, a specific CI inhibitor under clinical evaluation^56,57^ and Metformin

Hydrochloride, a treatment for diabetes that inhibits CI^58–61^. Using high content cellular imaging, we assessed impact on cell growth (by cell number) and death (by PI positivity) in HCT116 cells treated with 5FU alone, CI inhibitor alone or a combination of CI inhibitor and 5FU concurrently for 72 hrs. We observed a significant decrease in growth and increase in cell death in response to combination treatments with both IACS/5FU (Ext. data figure 7a) and Metformin/5FU (Ext. data figure 7b). We confirmed this using SRB staining (which stains total cellular protein, used as a surrogate for cell survival not dependent on ATP activity, Ext data figure 7c) after 72 hr treatment, and colony formation assays (Ext. data figure 7d, 72 hrs treatment, growth assessed after 10 days). Metformin/5FU treatment induced higher levels of cell death than IACS/5FU, and recent clinical trial data suggest neurotoxicity issues with IACS^57^, therefore we moved forward with 5FU/Metformin for *in vivo* evaluation.

We first assessed efficacy in HCT116 cell subcutaneous xenograft models. Tumours grown in athymic nude mice were randomised when average tumour value reached 100 mm^3^ and allocated to one of four treatment groups: 1: Saline control; 2: 5FU (10 mg/kg/day); 3: Metformin (250 mg/kg/day) and 4: 5FU/Metformin. Animals were treated daily for one week and change in tumour volume normalised to Day 0. 5FU resulted in modest, non-significant decrease in tumour growth compared to control, and Metformin alone had no significant impact on tumour control. However, the combination of 5FU with Metformin treatment showed significant growth inhibition from day 5 of treatment (Figure 6a). Tumours were sampled and FFPE sections stained for TOM20 expression by immunohistochemistry (IHC). TOM20 intensity was quantified here by imageJ. A significant increase in TOM20 in 5FU treated tumours was evident (Figure 6b), however, this was blunted by the 5FU with Metformin combination will be clinically effective.

**Figure 6:**
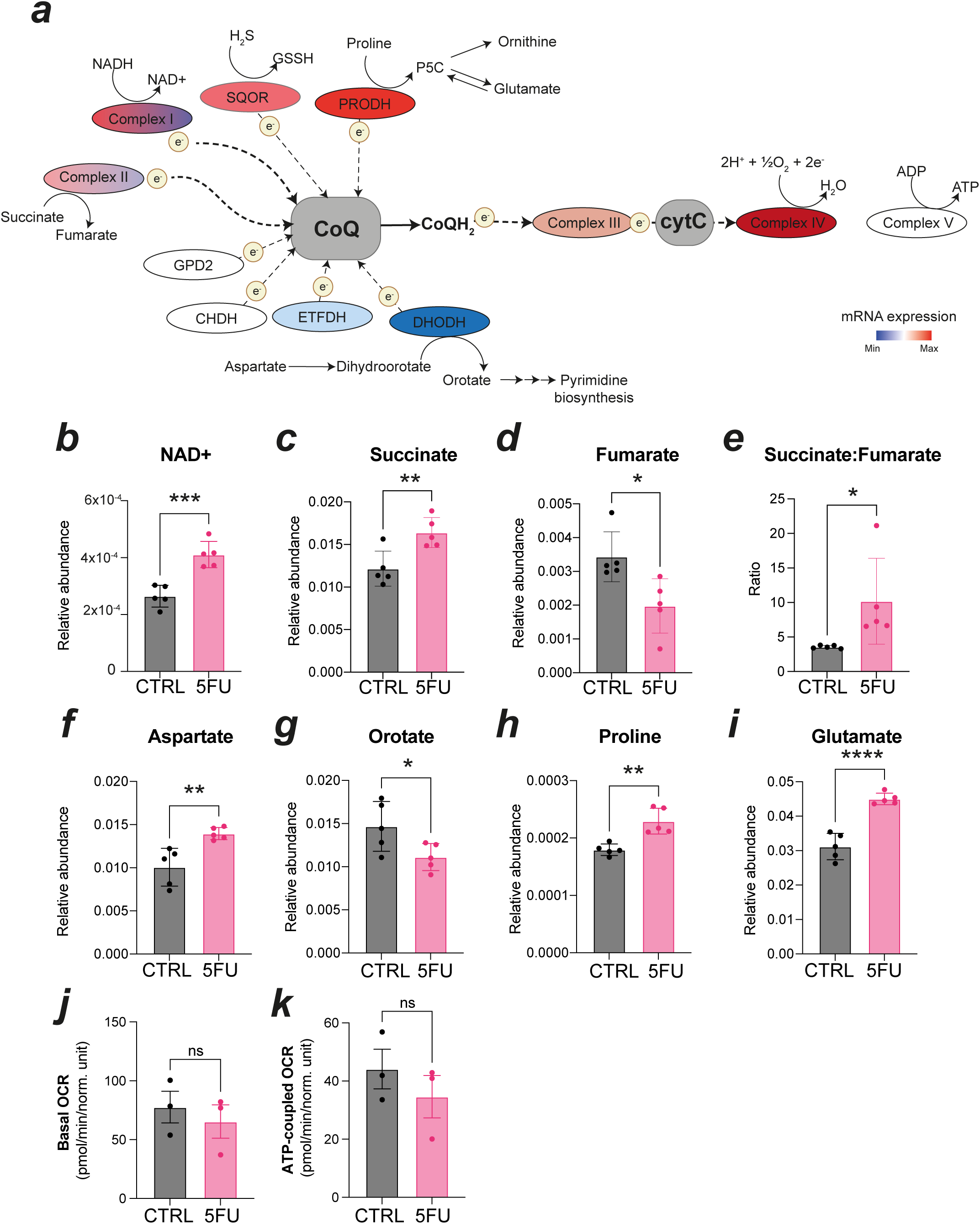
Inhibition of mitochondrial metabolism by metformin treatment increases anti-cancer response to 5FU. (a) HCT116 subcutaneous xenograft growth over time in mice treated with PBS (CTRL), 5FU, Metformin or 5FU & Metformin combination, with tumour growth normalised to day zero. Data points show mean tumour volume per group ± s.e.m. at each timepoint (n=4/5 animals per group); (b) representative TOM20 IHC staining of tumours collected at treatment endpoint, with quantification of intensity following Image J analysis. Data shows staining intensity per tumour ± s.e.m. Scale shown = 50 µm; Schematic representing experimental design for tumour induction and 5FU/Metformin treatment in *A*(Het)*KP* (c) and *AKP* (e) mice; Representative images and quantification of cleaved caspase 3 staining (% positive/total cells) by IHC in colon lesions (d, from c) and tumours (f, from e). Positive cells and clusters indicated by red arrows. Scale shown = 50 µm. Data points show mean % cleaved caspase 3 positive cells/mouse (d) or % cleaved caspase 3 positive cells/tumour (f), ± s.e.m.; (g) schematic of experimental design of 5FU or 5FU & Metformin treatment in *AKP* mice for survival analysis; (h) survival outcomes by Kaplan Meier plot, analysed using log-rank test, with animal number per cohort and P value reported. ****p<0.0001, ***p<0.001, **p<0.01, *p<0.05, ns = not significant, determined by two-way ANOVA with Šídák’s multiple comparisons test (a) or one-way ANOVA with Dunnett’s multiple comparisons test against CTRL samples (b, d, f).

**Figure 7:**
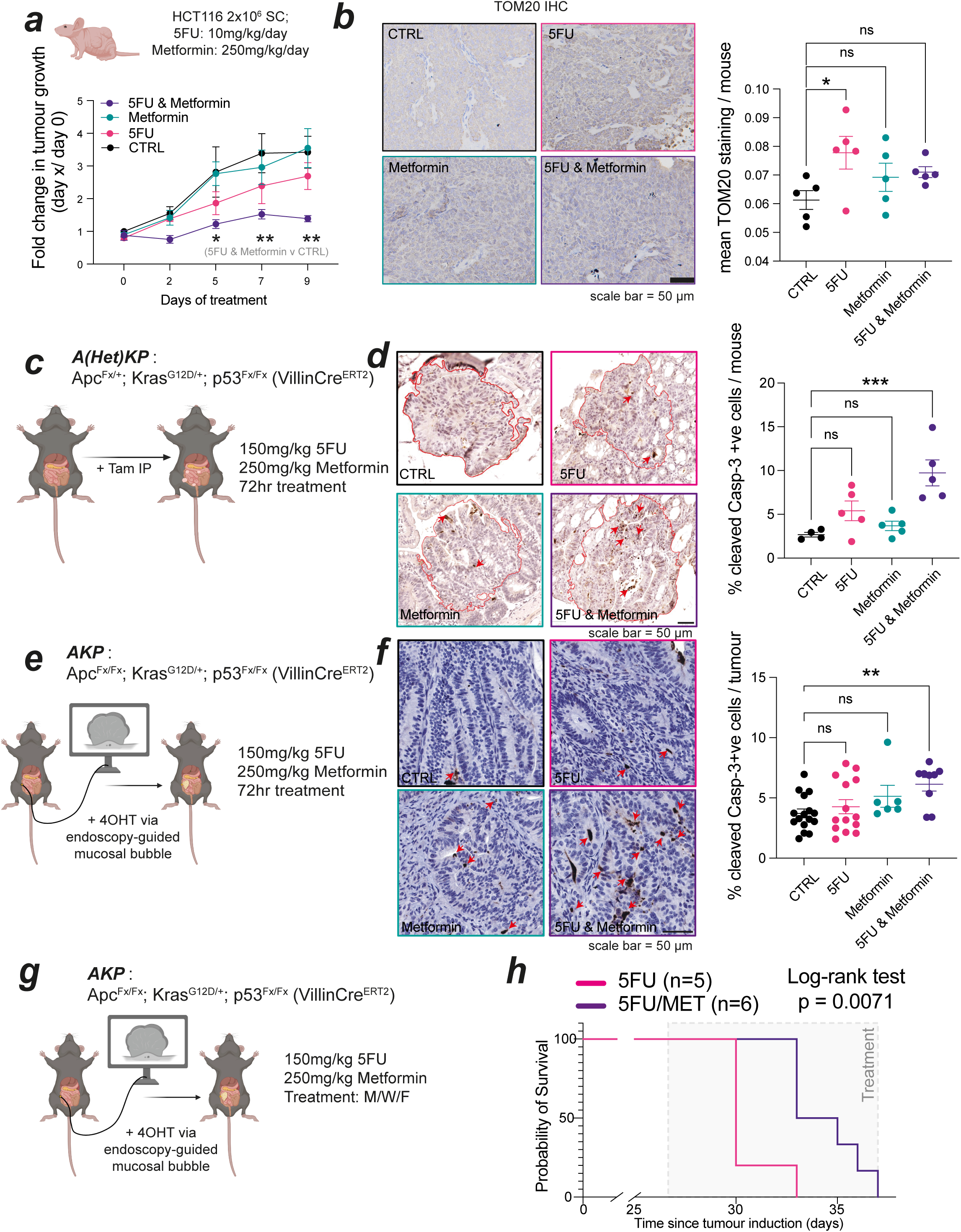
Inhibition of mitochondrial metabolism by metformin treatment increases anti-cancer response to 5FU.

We then evaluated the efficacy of this treatment combination in the orthotopic, immunocompetent models of CRC described above. We tested the sensitivity of *AKP* organoids isolated from these models to 5FU, Metformin or 5FU with Metformin combination treatment and demonstrate a significant increase in growth arrest with the combination when compared to single treatments alone (Ext. data figure 7e). Subsequently, *VilCre;Apc^Fx/+^;Kras^G12D/+^;p53^Fx/Fx^*mice were injected with 80 mg/kg tamoxifen i.p., and low - grade tumours developed over 3-6 weeks in the small and large intestines. At onset of tumour symptoms, animals were randomised and treated with 150 mg/kg 5FU (i.p.), 250 mg/kg Metformin (o.g.), a combination of both, and/or their retrospective saline vehicles, and tumour tissue collected after 72 hrs treatment (Figure 6c). FFPE sections were stained for cleaved caspase 3, a marker of apoptotic cell death, and percentage positive cells per tumour quantified by QuPath^62^. Multiple large intestine tumours were assessed per animal, and the average % cleaved caspase 3 positive cells per tumour per mouse is shown. Only in the combination group was there significant induction of cell death compared to control groups (Figure 6d). To assess the efficacy of this treatment combination in more aggressive colon tumours, we then used *VilCre;Apc^Fx/Fx^;Kras^G12D/+^;p53^Fx/Fx^*mice and induced tumours specifically in the colon tissue of mice as described above (Figure 2). First, tumour bearing animals were treated for 72 hrs (Figure 6e), and FFPE tumour sections analysed for cleaved caspase 3. As was observed in Figure 6d, combination treatment also increases the % apoptosis induced in these more advanced colon tumours relative to tumour animals treated with vehicle control (Figure 6f). Lastly, we tested if this 5FU with Metformin combination treatment could extend survival when compared to 5FU alone. Tumours were induced and upon presentation of clinical symptoms of tumour burden, animals were randomised and treated with 150 mg/kg 5FU by i.p. injection three times per week, with 250 mg/kg Metformin or vehicle control delivered concurrently by o.g. (Figure 6g). Animals began treatment ∼26 days post-induction and survival recorded as days post tumour induction at which clinical endpoint was reached. Kaplan-Meier survival analysis demonstrates that 5FU with Metformin treatment resulted in a significant extension in survival (Figure 6h, Log-rank test, p = 0.0071), and a Hazard Ratio of 5FU with Metformin/5FU alone of 0.3330.

Taken together, these *in vivo* data from multiple CRC models support our conclusion that inhibition of CI following Metformin treatment can enhance the response of 5FU treatment by increasing tumour cell death and impeding tumour growth.

## Summary

Drug resistance is not a new issue but remains the one that poses greatest difficulty for effective treatment of cancer patients worldwide^63^. As elegantly described by Vasan and colleagues, the combination of drugs targeting distinct, non-overlapping pathways has proven effective in enhancing treatment efficacy. In CRC, combining 5FU with DNA damaging agents Oxaliplatin and/or Irinotecan was an empirical choice rather than a rationally designed strategy, and no new combination partners have been added since the introduction of the biological EGFR and VEGF-targeting agents in the early 2000s. As fluoropyrimidine-based therapy will remain a major treatment modality for advanced CRC patients for the foreseeable future, the need for better, more effective chemotherapy combinations in biologically-selected populations to overcome drug resistance is an urgent, unmet need for CRC patients worldwide.

Our findings presented herein describe a novel adaptation adopted by CRC cells in response to 5FU-based standard-of-care treatment that results in a targetable vulnerability. By increasing mitochondrial biogenesis, signalling and activity, 5FU-treated CRC cells maintain the processes required for cell growth under stress conditions, and the functional inhibition of this adaptive response greatly increased the effectiveness of 5FU across human and murine CRC models, both *in vitro* and *in vivo*.

Here, we present a comprehensive, systematic analysis of the contribution of mitochondrial metabolism to chemotherapy persistence and resistance in CRC using 2D and 3D cultures, *in in vivo* models and by interrogating clinical data. Upregulation of oxidative mitochondrial metabolism is an accepted method of resistance to other therapeutic interventions in various diseases such as pancreatic cancer and AML^64,65^. Our findings are in line with this and demonstrate that metabolic reprogramming is a key driver of drug resistance in CRC. Moreover, since the inception of this study, multiple metabolic mechanisms promoting drug resistance have been suggested for 5FU treatment, including changes in amino acid metabolism^66,67^, serine/glycine metabolism^68–70^ and glycolysis^71,72^, corroborated within our study and independently validating our findings. For the first time we now demonstrate that oxidative mitochondrial signalling can be used as a biomarker of response to FP-based chemotherapy treatment in colorectal patients and could provide clinical utility in stratifying higher risk patients that would benefit from metabolically targeted combination therapy approaches.

Inhibition of oxidative metabolism to target cancer cell proliferation using various inhibitors has shown promise in both human and murine preclinical models^73^. Our choice of metformin as a proof of concept test *in vivo* was based on a number of factors: 1. CI inhibitors preformed best in testing of an ETC targeted drug panel (data not presented here given spatial constraints); 2. It could be taken forward into *in vivo* studies; 3. The specific CI inhibitor IACS-010759 was recently reported to promote neurotoxicity making this an unattractive agent for combination therapy in CRC given the neurotoxicity long-recognised with oxaliplatin treatment^57^; 4. Another candidate from the *in vitro* screen, Phenformin has been linked to increased lactic acidosis in patients^74^; 5. Metformin is safe, well tolerated and cost-effective, making it an attractive addition to a cytotoxic therapeutic regimen, and can be administered orally alongside infusional 5FU. Metformin treatment has been previously explored in CRC; however contradictory findings have been reported^75–77^. Retrospective studies suggest a beneficial effect on CRC risk and recurrence in diabetic patients, but the action of these drugs on tumours in diabetic patients could be quite different from non-diabetic patients given differences in central metabolic programmes and confounding effects on overall mortality. Furthermore, heterogeneity in tumour stage (or indeed location), patient populations and treatment dose and duration is likely to confound reported response data, particularly given the links between changes in oxidative metabolism and metastatic spread^78–81^. While a small number of randomised trials investigating metformin use in CRC patients are ongoing, these are in unselected populations and may also include other interventions (such as aspirin and physical activity). Therefore, clinical testing in controlled populations is ultimately required to assess if this preclinical efficacy is also observed in CRC patients with accompanying translational research to confirm our proposed underpinning mechanisms of action. As more specific CI-targeted drugs enter clinical pipelines, these can also be deployed in defined CRC cohorts to examine combination efficacy.

Our data demonstrate that the adaptive metabolic response to 5FU is mediated through the inhibition of TS function, as we recapitulate these phenotypes by genetic silencing or targeted inhibition of TS using antifolates. Over 5 million patients are treated with TS inhibitors annually^fn^ ^82,83^, expanding our findings beyond the CRC field to other disease indications that use Pemetrexed or Methotrexate, such as lung cancer. Indeed, we found a similar mitochondrial response to Pemetrexed treatment in three independent KRAS mutant non-small cell lung cancer cell lines (Ext. data figure 3c). This highlights not only the key role that mitochondrial metabolism plays in modulating response and resistance to TS inhibitors, but the wider opportunity afforded to enhance TS inhibitor mediated anti-cancer effects by targeting mitochondrial metabolism using Metformin or indeed other compounds with similar mitochondrial impacts.

In summary, we present a panoptic analysis of the role of mitochondrial metabolism in driving resistance to 5FU, the most commonly used chemotherapy worldwide. We show that the heterogeneity in metabolic signalling observed across CRC samples could explain, and be used to predict, poor clinical responses in patients with locally advanced CRC. We demonstrate applicability beyond 5FU treatment in CRC to any TS targeted therapeutics in other disease subtypes. Finally, we demonstrate the potential of rationally designing targeted combination therapy regimens for CRC, and with the compelling preclinical data presented herein, a clinical trial demonstrating efficacy and mechanism of action in patients with CRC is in development.

## Figure legends

**Extended Data figure 1:**
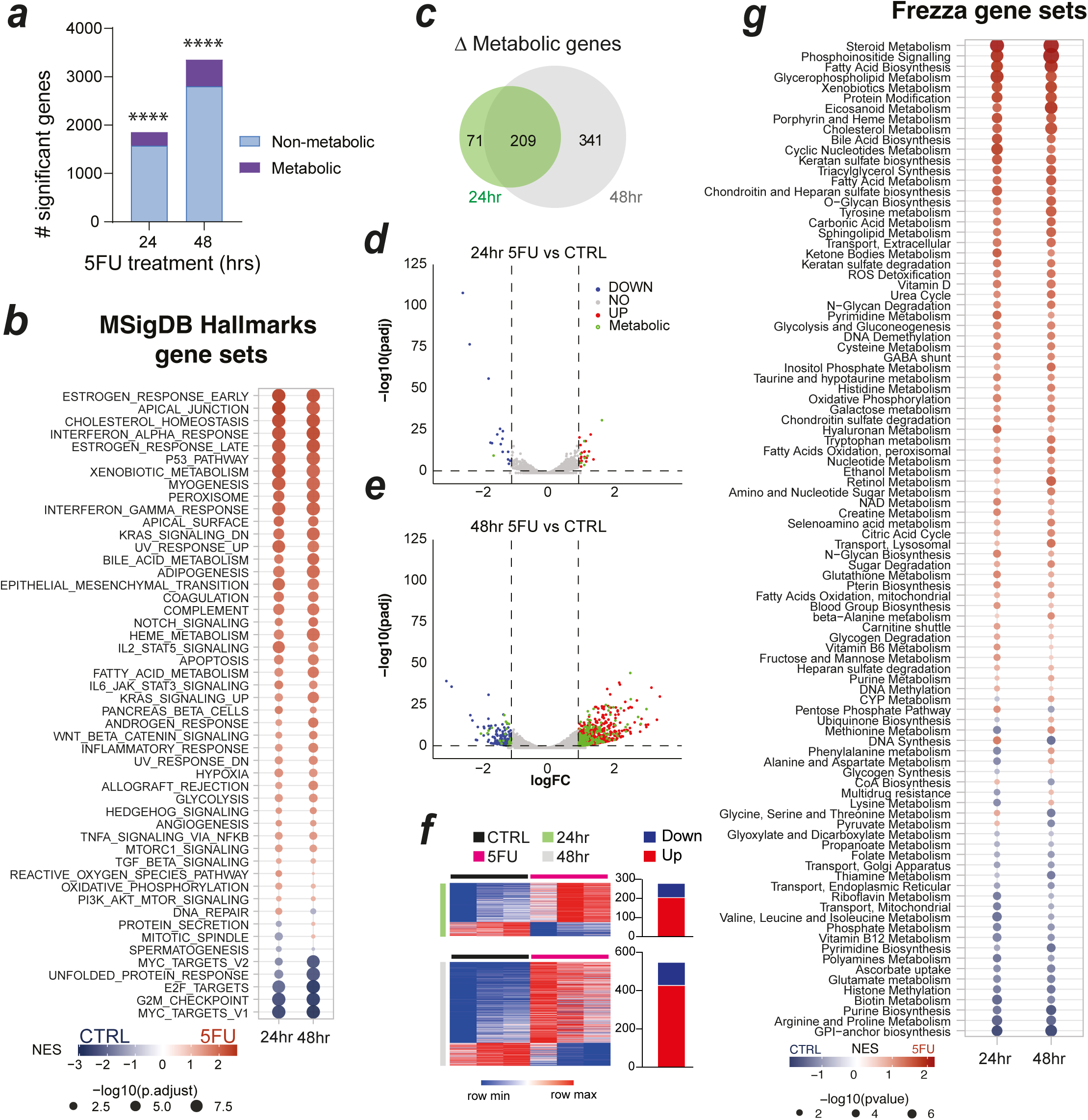
5FU reprograms metabolic signalling networks. (a) Number of significant genes (defined as adj. P value <0.05) following DESeq2 analysis of transcriptional data (RNASeq) from HCT116 cells treated for 24 and 48 hr with 5FU. Metabolic genes (defined in ^10,19^) are indicated, with over-representation significance (****P<0.0001) calculated by Fisher’s exact test; (b) GSEA analysis of data from (a) run against MSigDB Hallmarks genesets; (c) Metabolic genes from (a) at 24 and 48 hrs; Volcano plots of differentially expressed genes at 24 (d) and 48 (e) hrs, with metabolic genes highlighted (green); Heatmaps and summary bar charts of metabolic gene expression patterns of genes from (c); (g) GSEA analysis of data from (d & e) run against metabolic gene sets (defined in ^10^).

**Extended data figure 2:**
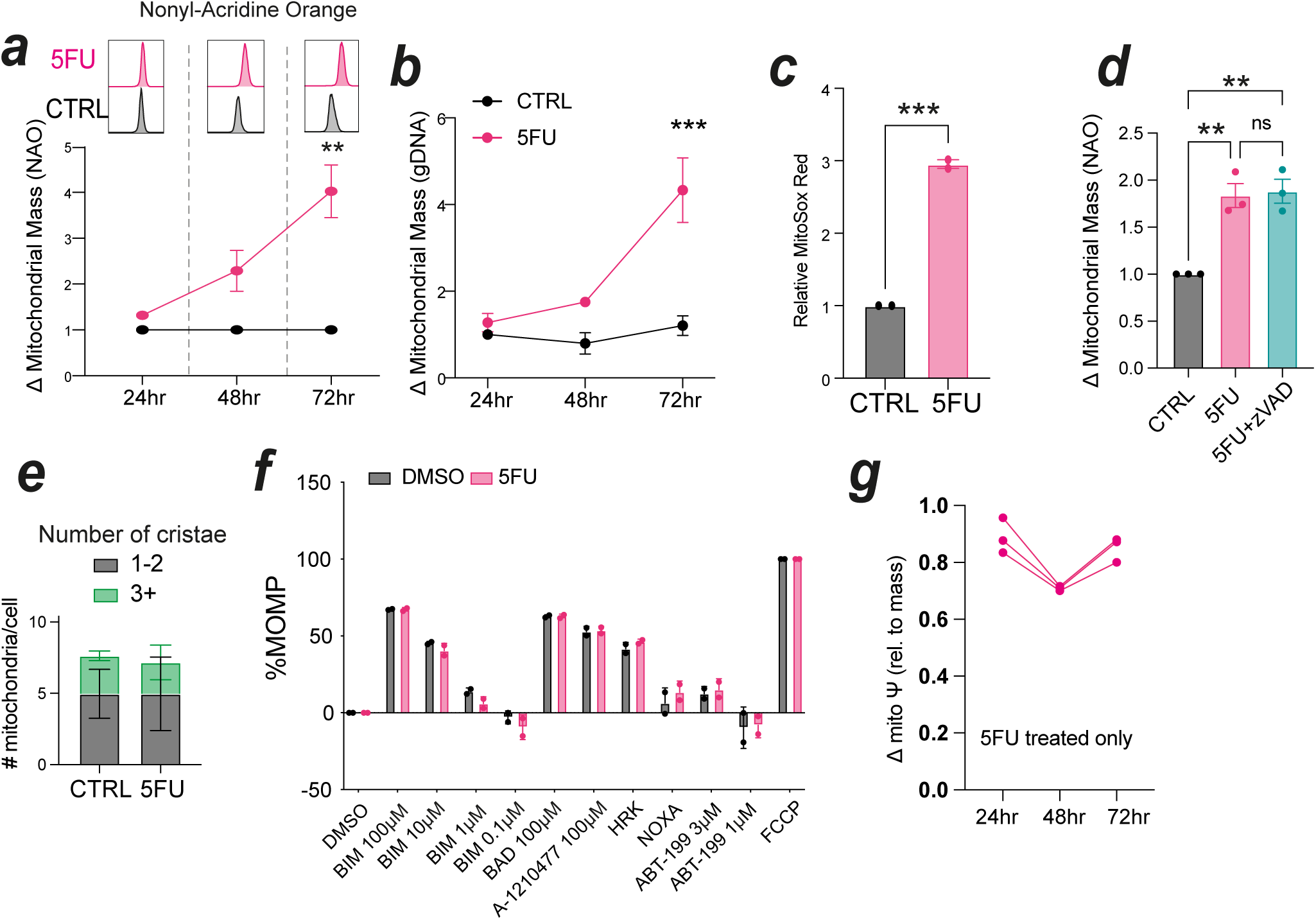
5FU treatment increased mitochondrial mass and activity independent of apoptosis. Change in mitochondrial mass in HCT116 cells treated ± 5FU over 72 hrs measured by Nonyl-Acridine Orange (NAO) staining (a) or mitochondrial DNA content (b), data point shows biological mean, ± s.e.m. of n=2 experiments; (c) Mitochondrial ROS in HCT116 cells ± 72 hrs 5FU measured by MitoSox Red staining. Data shows two independent biological means calculated from technical triplicates, with mean ± s.e.m.; Change in mitochondrial mass measured by NAO staining of HCT116 cells treated ± 5FU or 5FU + z-VAD-fmk (d). Data points show 3 biological replicates ± s.e.m; (e) graph illustrating number of mitochondria analysed per cell, with cristae number noted, ± s.d.; (f) Degree of mitochondrial outer membrane permeabilization (% MOMP) in HCT116 cells ± 5FU when treated with compounds as indicated. Data shows technical replicates ± s.d.; (g) Change in mitochondrial membrane potential, calculated by degree of TMRE staining, relative to mitochondrial mass (a) at each timepoint for HCT116 cells treated with 5FU. ****p<0.0001, ***p<0.001, **p<0.01, *p<0.05, calculated by two-way ANOVA with Šídák’s multiple comparisons test (a,b), t-test (c), one-way ANOVA with Dunnett’s multiple comparisons test (d).

**Extended data figure 3:**
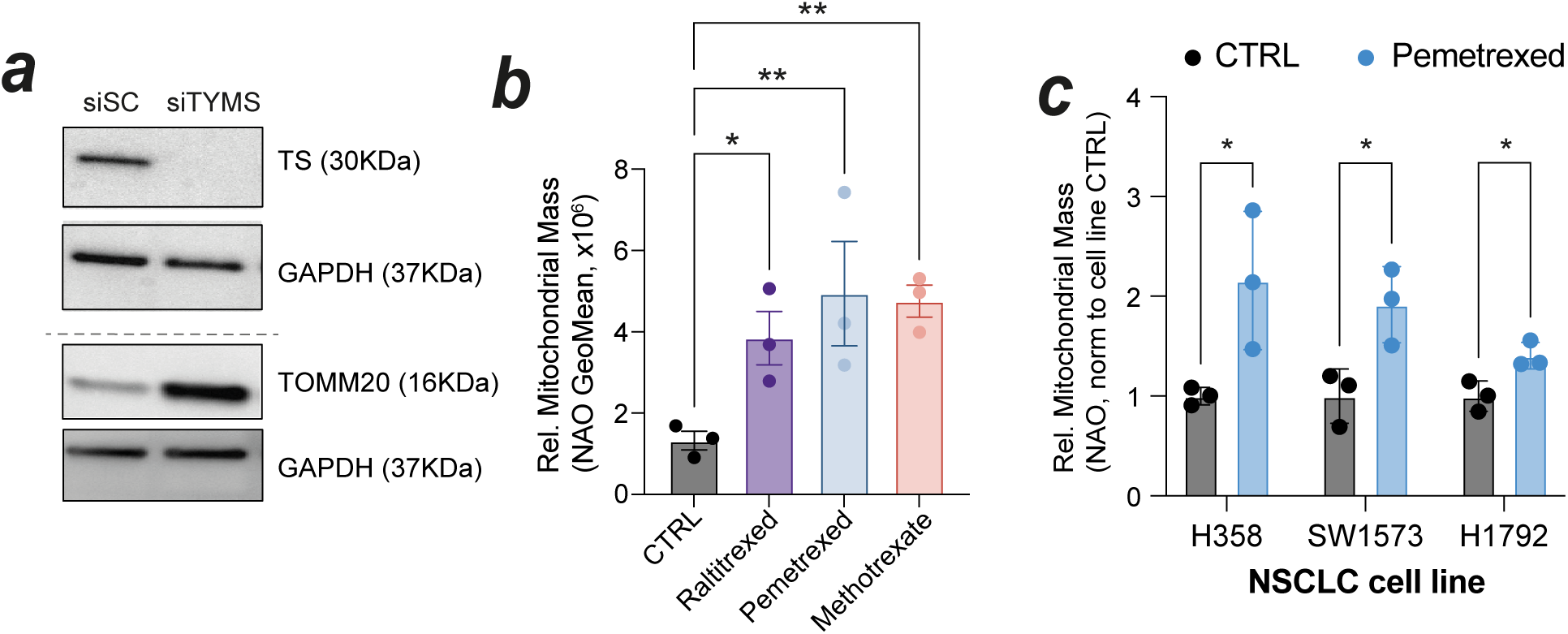
Thymidylate Synthase inhibition drives increased mitochondrial mass. (a) Western blotting analysis of protein lysates from HCT116 cells transfected for 72 hrs with control (CTRL) or TYMS siRNA; Mitochondrial mass assessed by NAO staining in HCT116 cells treated with TS-targeting therapies (b) or non-small cell lung cancer (NSCLC) cell lines treated with pemetrexed for 72 hrs. Data shows biological (b) or technical (c) replicates, with mean ± s.e.m. (b) or s.d. (c). **p<0.01, *p<0.05, by one-way ANOVA with Dunnett’s multiple comparisons test (b) or paired t-test (c).

**Extended data figure 4:**
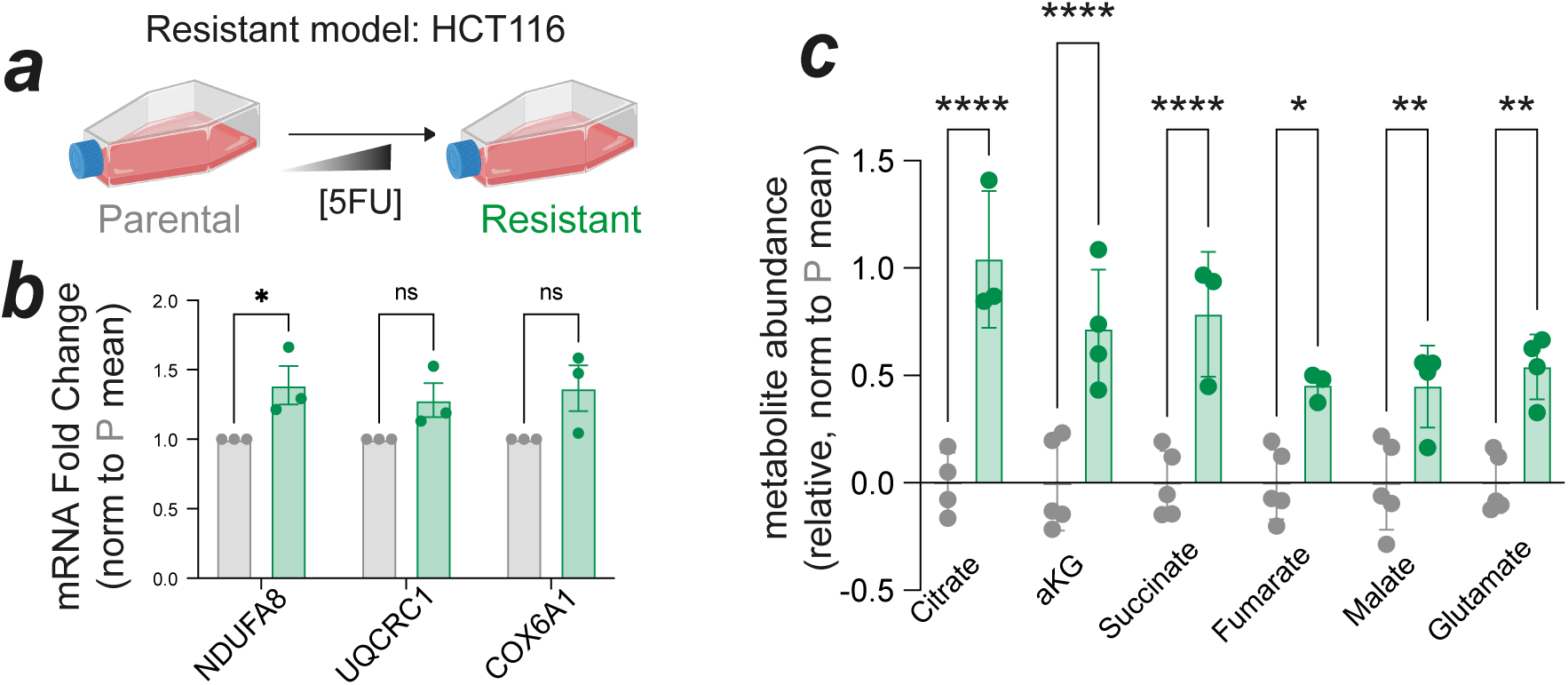
5FU-resistant HCT116 cells have increased mitochondrial metabolism. (a) Schematic of generation of resistant HCT116 cells by chronic culture methods as previously reported ^52^; (b) relative expression of electron transport chain (ETC) genes by qPCR analysis, relative to 18S and normalised to paired Parental sample. Data shows biological replicates ± s.e.m.; (c) relative mitochondrial metabolite abundance quantified by LC-MS, relative to total ion count/cell, normalised to parental mean. Data shows technical replicates ± s.d.; ****p<0.0001, ***p<0.001, **p<0.01, *p<0.05 by T-test (b) and two-way ANOVA with Šídák’s multiple comparisons test.

**Extended data figure 5:**
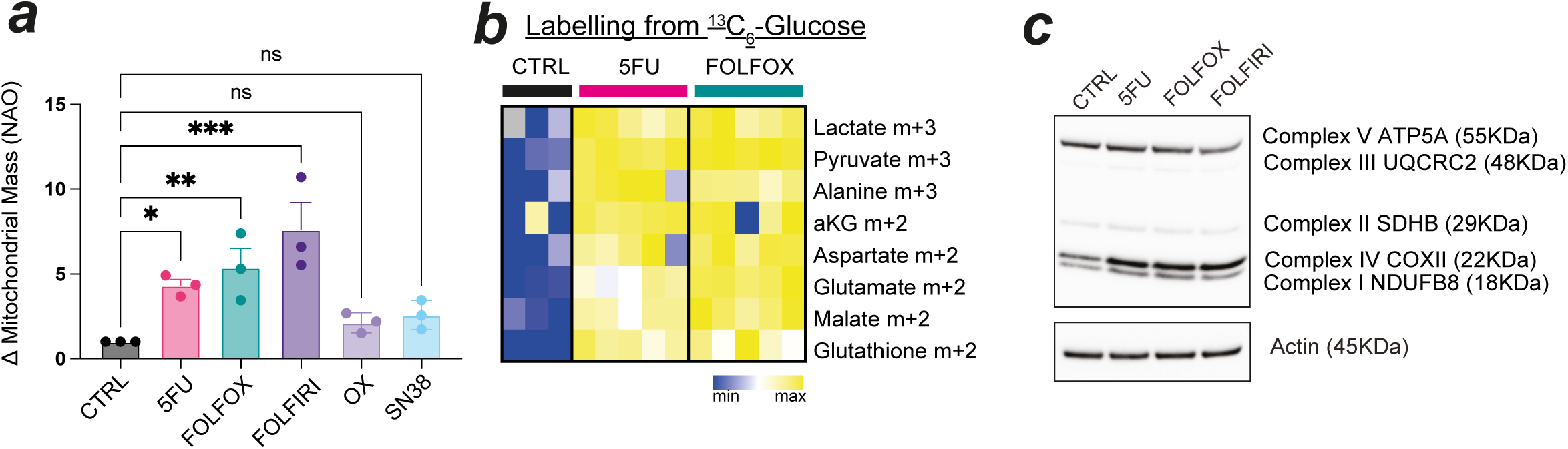
Mitochondrial phenotypes are conserved upon combination chemotherapy treatment. Relative mitochondrial mass measured by NAO staining in HCT116 cells treated with 5FU, 1μM Oxaliplatin, 5nM SN38, FOLFOX (5FU + 1μM Oxaliplatin) or FOLFIRI (5FU + 5nM SN38). Data shows biological replicates, normalised to control (CTRL) ± s.e.m.; (b) Heatmap shows relative abundance of labelled metabolite isotopologues indicated, derived from ^13^C_6_-Glucose, in HCT116 cells treated with 5FU or FOLFOX against control (CTRL) samples. Technical replicates are shown, data representative of two independent experiments; (c) Western blotting analysis of protein lysates derived from HCT116 cells treated with 5FU, FOLFOX or FOLFIRI against CTRL samples. ***p<0.001, **p<0.01, *p<0.05, ns = not significant, determined by one-way ANOVA with Dunnett’s multiple comparisons test.

**Extended data figure 6:**
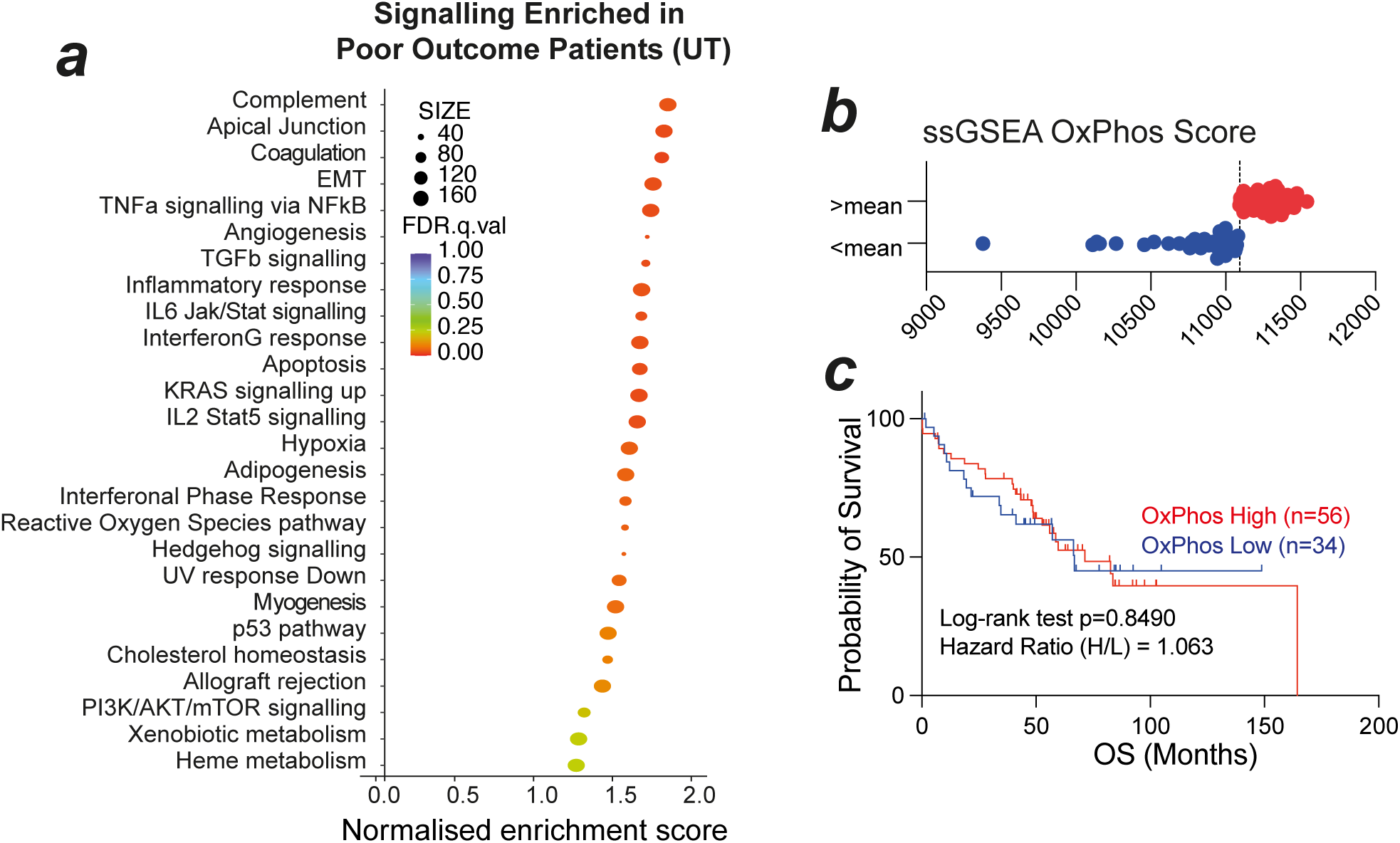
Mitochondrial metabolism does not predict poor outcomes following surgery alone. Analyses of patients included in Taxonomy cohort that did not receive adjuvant 5FU-based chemotherapy treatment; (a) GSEA analysis of upper vs lower tertiles ranked on overall survival, graph shows Normalised Enrichment Score (NES) for Poor Outcomes patients; (b) OxPhos scores per patient, split by cohort mean; (c) Kaplan Meier Overall Survival (OS) of patients split as in (b), with Log rank test and Hazard Ratio analyses reported.

**Extended data figure 7:**
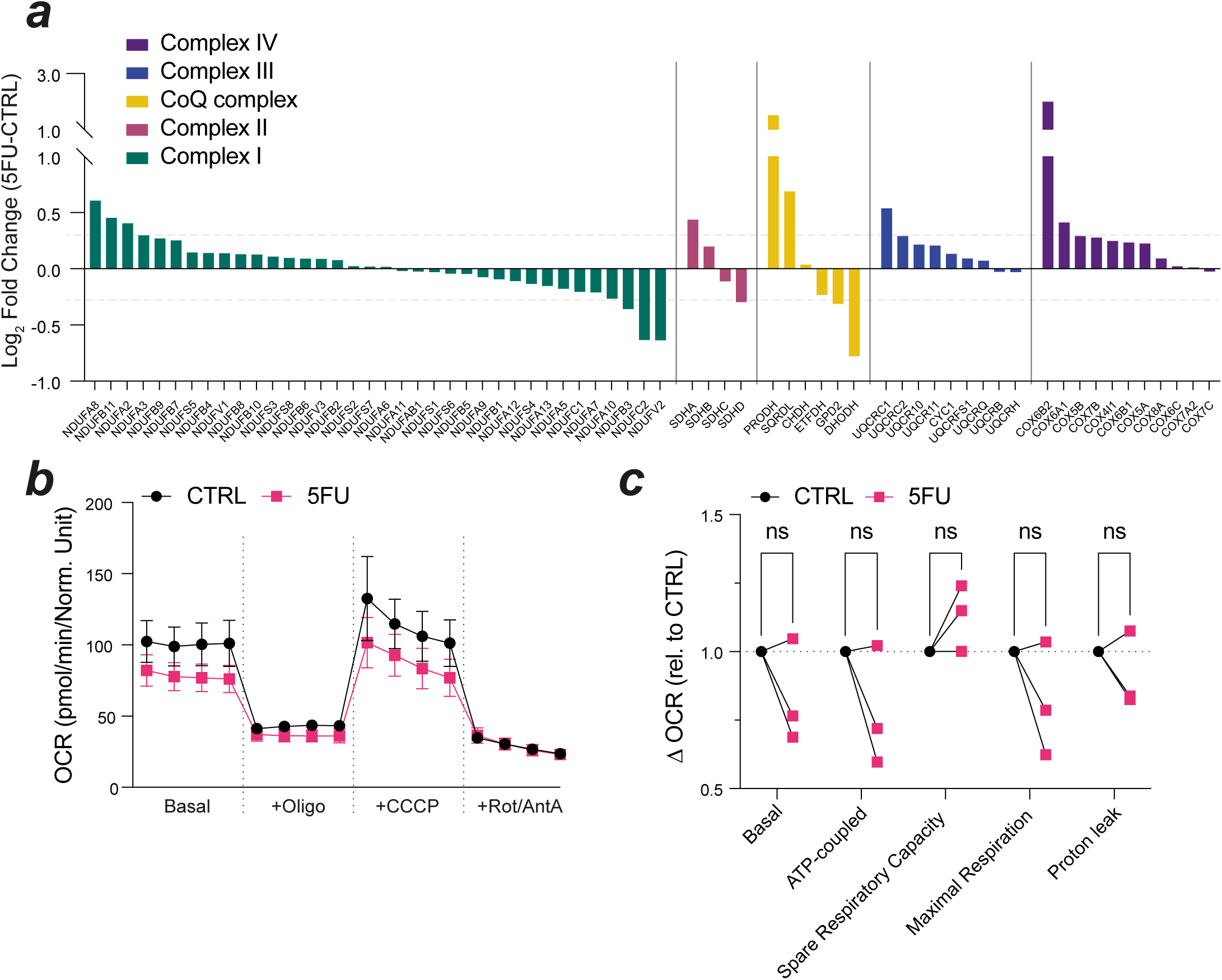
Inhibition of mitochondrial CI increases 5FU efficacy. High content images (magnification 10x) from HCT116 cells stained with Hoescht (blue) and Propidium Iodide (green) following treatment ± 5FU, ± CI inhibitors IACS-010759 (IACS, a) or Metformin Hydrochloride (MET, b) for 72 hrs. Change in Growth (number of Hoescht +ve cells relative to vehicle control and media only control) and Death (% PI positive cells relative to vehicle control) is shown, data points show technical replicates ± s.d.; with mean Growth Arrest (1-Growth FOC) and mean %PI positive ± s.d. denoted; (c) HCT116 cells treated ± 5FU ± IACS-010759/Metformin for 72 hrs, with change in growth calculated by SRB staining and data normalised to CTRL. Heatmaps show mean fraction of control; (d) representative images (magnification 10x) with quantification of n=3 biological replicates ± s.e.m. of colony formation assay in HCT116 cells treated as indicated; (e) *AKP* organoids treated as indicated with 5 μM 5FU and/or 3 mM Metformin (MET) for 72 hrs. Representative images (magnification 4x) and quantification of relative organoid size shown. Data points show mean of independent organoid cultures, ± s.e.m.; ****p<0.0001, ***p<0.001, **p<0.01, *p<0.05, ns = not significant, determined by one-way ANOVA with Dunnett’s multiple comparisons test.

**Extended data figure 8:**
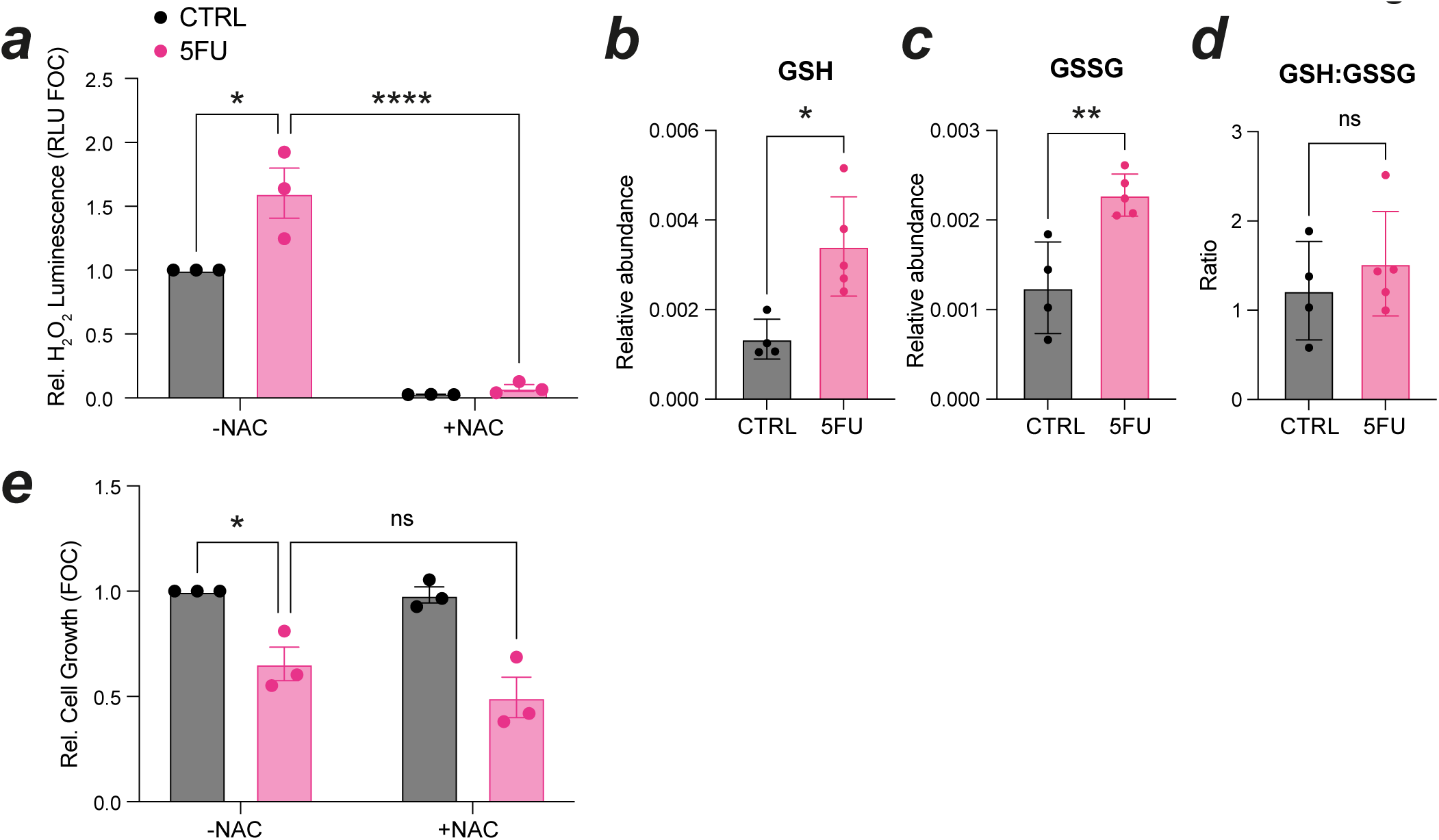
Increased ROS buffering does not rescue 5FU-induced growth arrest.

**Extended data figure 9:**
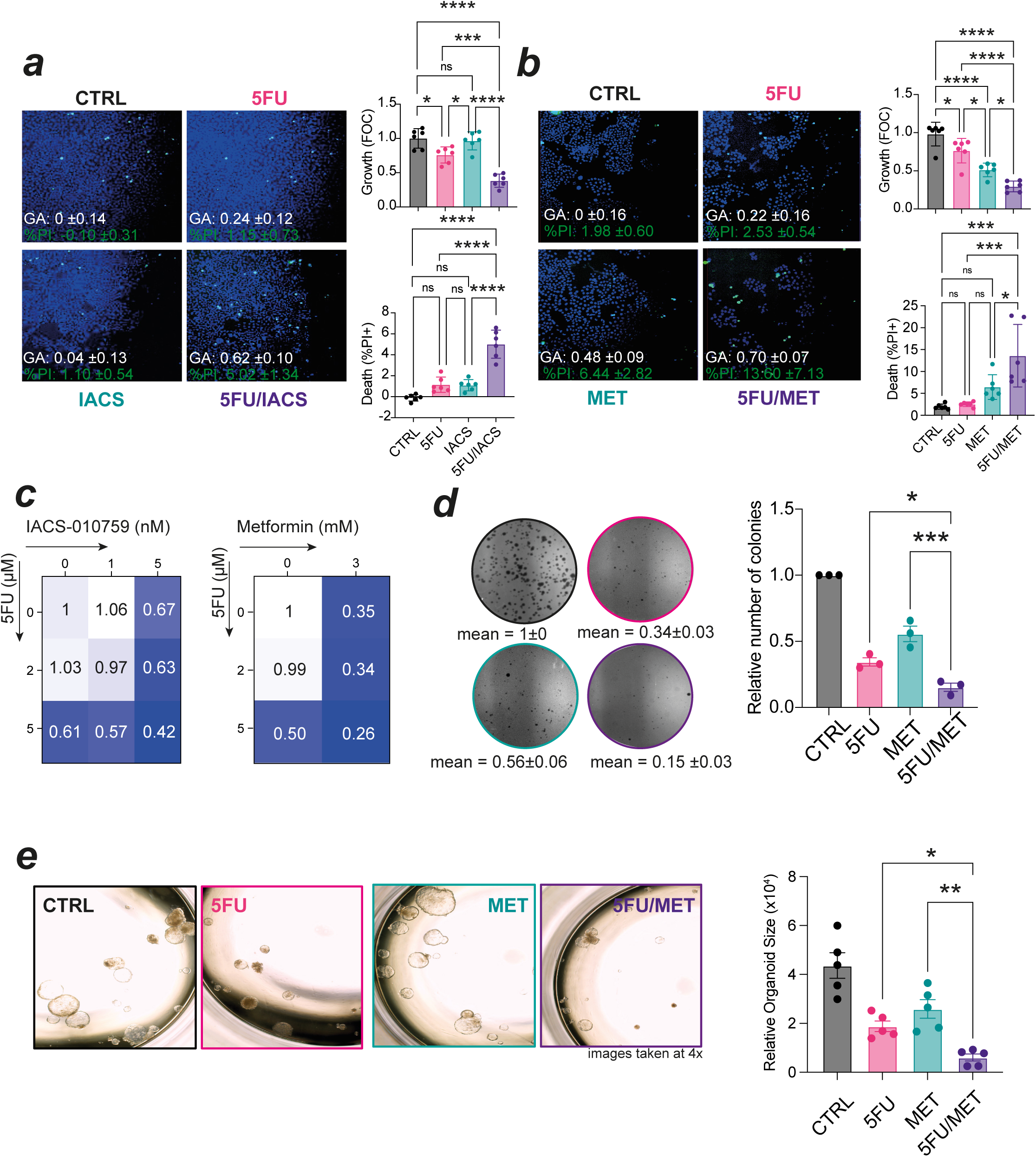
Inhibition of mitochondrial CI increases 5FU efficacy.

## Methods

### Cell culture

HCT116 CRC cell line was obtained from ATCC and maintained in DMEM with 10% dialysed foetal bovine serum (dFBS), 100 U/ml penicillin and 100 μg/ml streptomycin (ThermoFisher, 15070063), 2 mM L-glutamine (ThermoFisher, 25030081) and 1 mM sodium pyruvate (ThermoFisher, 11360070). Cell lines were maintained at 37 °C in 5% CO_2_ humidified incubators, and routinely tested for mycoplasma using MycoAlert^TM^ Kit (Lonza). 5FU resistant HCT116 cells were generated by chronic exposure to 5FU. Glucose/Galactose adapted cells were cultured in DMEM containing 10% dFBS, 100 U/ml penicillin and 100 μg/ml streptomycin, 1 mM sodium pyruvate and 25 mM glucose (Sigma-Aldrich, G7021-1KG) or 25 mM galactose (Sigma-Aldrich, G5388-500G) for a minimum of a month before being used experimentally. siRNA transfections using Lipofectamine^2000^ and ON-TARGETplus Human smart pools from Dharmacon against TYMS and a Scrambled Control pool according to manufacturer’s instructions.

### Organoid culture

Organoids were resuspended in Matrigel (Corning, 356237), plated in 6-well plates and supplemented with Advanced DMEM/F-12 medium (ThermoFisher, 11320033) with 2 mM L-glutamine, 10 mM HEPES (Sigma-Aldrich, H3375-25g), 100 U/ml penicillin and 100 μg/ml streptomycin, 1× B27 (ThermoFisher, 12587-010), 1x N-2 (ThermoFisher, 17502-048), 50 ng/ml EGF (Peprotech; AF-100-15) and 100 ng/ml Noggin (Peprotech, #250-38). 5FU resistant organoids were generated through chronic exposure to 5FU. To evaluate the effect of CI inhibition on growth, organoids were seeded as fragments in Matrigel and treated the following day with water (CTRL), 5 μM 5FU, 3 mM Metformin (MET) or in combination for 72 hr. Organoid diameter was measured using Image J and average calculated.

### Cell reagents

Cells/Organoids were treated with the following compounds as indicated: 5FU ( Abcam, ab142387-5g, 2-5 µM), Metformin (Cayman Chemicals, #13118, 3-5 mM), IACS-10759 (Selleckchem, S8731, 1-10 nM), zVAD-fmk (Sigma-Aldrich, 627610, 10 µM), siTYMS and siSC siRNA SmartPools (L- 004717-00-0005, Horizon Discovery, 10 nM), Oxaliplatin (Belfast City Hospital, 1 µM), SN38 (Selleckchem, S4908, 5 nM), Methotrexate (Merck, M1000000, Batch 8.0, 15 nM); Pemetrexed (Merck, PHR1596, Lot LRAC1932, 0.5 µM); Raltitrexed (Selleckchem, S1192, Lot S119204, 5 nM).

### Cell Viability Assay

Cell viability was measured using sulforhodamine B (SRB, Sigma-Aldrich, S1402-5G)^84^ and clonogenic assays. For the SRB assay, cells were seeded (5000 cells/well) in a 96-well plate and then treated with indicated drugs. At the experimental endpoint, the cells were fixed with 10% trichloroacetic acid (Sigma-Aldrich, T6399-500G) and stained with 0.05% SRB for 1 hr and then washed repeatedly with 1% acetic acid. The protein-bound dye was dissolved in 10 mM Tris base solution (ThermoFisher, 15504-020) to obtain a reading at 510nm. For the clonogenic assay, cells were seeded (5×10^2^ cells/well) in 6-well plates and treated with indicated drugs. Media was refreshed with drug-free media every 3 days and the cells were fixed after 10 days, stained with SRB and counted using CFU Scope (v1.6.0). The effect on cell growth was assessed as a percentage of cell viability, with vehicle-treated cells considered 100% viable. The IC_50_ of each drug was calculated using GraphPad software (GraphPad Software Inc.).

### Mitochondrial profiling

Cells were stained with Acridine Orange 10-nonyl bromide (NAO; ThermoFisher, A1372), Tetramethylrhodamine, Ethyl Ester Perchlorate (TMRE; Sigma-Aldrich, 87917) and MitoSOX Red (ThermoFisher, M36008). NAO (200 nM) or TMRE (25 nM) were added to the culture medium and incubated for 1 hr at 37 °C in 5% CO_2_. MitoSOX Red (5 µM) was added to the medium and incubated for 30 min at 37 °C in 5% CO_2_. Cells were washed, collected and resuspended in 0.3 ml of PBS. Cells were analysed by flow cytometry using a BD LSR II instrument. Cells were gated for singlets and the GeoMean NAO, TMRE or MitoSOX Red fluorescence was evaluated using FlowJo software.

gDNA was isolated using buffer (100 mM Tris-HCl, 5 mM EDTA, 0.2% SDS, 200 mM NaCl + proteinase K) and precipitation with isopropanol. Real time PCR was conducted to quantify the mitochondrial DNA relative to nuclear DNA as previously reported. 250 ng of target DNA was used per reaction. Primers had the following sequences:

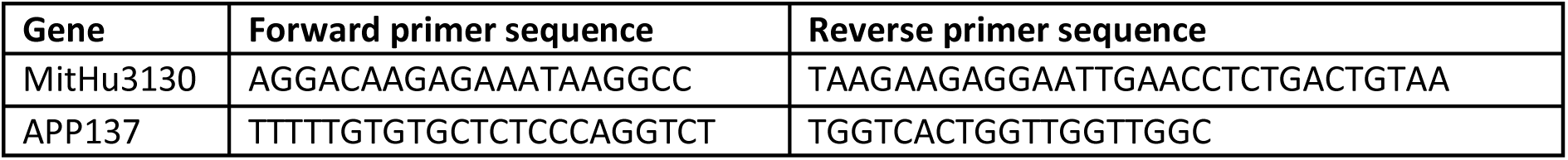

### CI analysis

The enzyme activity of CI was determined by a colorimetric analysis following the manufacturer’s protocol (Abcam, ab109721).

### Immunoblotting

Cells were washed with PBS and lysed with lysis buffer (150 mM NaCl, 20 mM EDTA, 50 mM Tris-HCl, 0.5% NP-40) containing protease inhibitors. Protein concentration was determined by BCA assay (ThermoFisher). After addition of Loading buffer, lysates were vortexed and supernatants were heated to 95 °C for 5 minutes. Then 30 μg of total protein were loaded on Novex^TM^ WedgeWell^TM^ Tris-Glycine gels (ThermoFisher, XP00120BOX) and blotted onto nitrocellulose Trans-Blot Turbo membrane (0.2 µm, Biorad, 170-4279) according to standard protocols. Membranes were incubated with the appropriate Primary antibody overnight at 4 °C. The following antibodies were used: Total OxPhos Human Antibody Cocktail (Abcam, ab110411; 1:500), rabbit anti-TOM20 (Abcam, ab186734; 1:1000), rabbit anti-OPA1 (Cell Signalling Technology, 80471; 1:1000), rabbit anti-DRP1 (Cell Signalling Technology, 8570; 1:1000), rabbit anti-Thymidylate Synthase (Cell Signalling Technology, 9045S; 1:1000), mouse anti-GAPDH (Santa Cruz Biotechnology, sc-47724; 1:10,000) and mouse anti-β-actin (Sigma-Aldrich, A5316; 1:10,000). Membranes were washed three times with PBS-T and incubated for 1 hr at room temperature with the corresponding horseradish peroxidase (HRP)-labelled Secondary antibodies: Goat Anti-Rabbit IgG (H+L) (Vector Laboratories, (BA-1000; 1:2000) and Rabbit Anti-Rat IgG (H+L) (Vector Laboratories, BA-4000; 1:2000). Protein expression was analysed and detected using the Western Lightning® Plus-ECL, Enhanced Chemiluminescence Substrate (PerkinElmer, NEL105001EA) and G:BOX ChemiXX6 gel doc system with GeneSys image software (Syngene). Densitometry completed using imageJ.

### Metabolomics analysis

1 ×10^5^ HCT116s treated ± 5 μM 5FU or FOLFOX (5FU + 1 μM Oxaliplatin) were supplemented with media containing uniformly labelled ^13^C-glucose (25 mM) for 6 hr before sampling. Metabolites were extracted from media (extracellular) and cell pellet (intracellular) in 50% MeOH: 30% Acetonitrile: 20% H_2_O + 100 ng/mL HEPES buffer (MEB). Samples were incubated at 4 °C for 15 minutes at 700 rpm (Thermomixer), before centrifugation at 15,700 rcf. Supernatant was transferred to vials and dried using an Eppendorf Concentrator Plus at 30 °C for up to 7 hr. Samples are reconstituted in 100 µl 50% acetonitrile (aq), before centrifugation at 2000 rpm. Supernatant was transferred to vials for mass spectrometry analysis. LC separatation was performed using an Agilent InfinityLab Poroshell 120 HILIC-Z (2.1 x 100 mm, 2.7 μm), PEEK-lined, column on an Agilent G7167B and G7120A multisampler and pump, respectively. Separation was achieved using a gradient of 10 mM ammonium acetate in water (mobile phase A; pH 9.0) and 10 mM ammonium acetate in 90% acetonitrile (aq) (mobile phase B (MPB); pH 9.0). Flow rate was maintained at 0.5 mL/minute and the gradient was as follows: 0.0 minutes 100% of MPB, 0.0-11.5 minutes 70% of MPB, 11.5-12.5 minutes 60% of MPB, 12.5-15.5 minutes 100% of MPB. Mass spectrometer (Agilent Dual ESI G6545B quadrupole time-of-flight (Q-ToF)) was operated in negative ion mode. Spectra were analysed using Agilent MassHunter Qualitative Analysis and Agilent MassHunter Profinder software by referencing to an internal library of compounds. Relative metabolite abundance was calculated as percentage of total metabolite pool.

### Transmission electron microscopy

HCT116 cells were treated with 5 µM 5FU for 72 hr, before being pelleted and fixed with 3% glutaraldehyde (Agar Scientific) in cacodylate (CACO). Cells were then washed three times for 10 minutes each in CACO buffer. Cells were post-fixed for 1 hr with 1% osmium tetroxide (Agar Scientific) in CACO buffer, dehydrated in a graded series of methanol and embedded in LR white resin. Samples are sectioned using an ultramicrotome (Leica EM UC6) to 60-90 nm, transferred to copper grids and left to dry before imaging. Electron micrographs were obtained using a transmission electron microscope (Hitachi) at x40,000 magnification. Analysis was carried out using ImageJ software. Images with less than 5 mitochondria were discounted. The number of cristae were manually counted. The aspect ratio was calculated as major axis/minor axis.

### BH3 profiling

HCT116 cells were treated with 5 µM 5-FU for 24 hours. BH3 profiling was then performed using whole cell (JC-1) plate-based fluorimetry. BH3 peptides/mimetics in DTEP Buffer (300 mM Trehalose, 10 mM HEPES-KOH, 0.1 % w/v BSA, 1 mM EDTA, 1 mM EGTA, 80 mM KCl, 5 mM succinate, final pH 7.4) were plated at 70 μM/L (unless otherwise stated) in triplicate in a black 384-well plate. The sequence of the BH3 peptides are listed previously^85,86^. HCT116 cell lines were harvested, washed and resuspended in DTEP buffer. An equal volume of dye Mastermix (1 μM JC-1, 0.005% digitonin, 10 μg/ul oligomycin, 5 mM B-mercaptoethanol in DTEP buffer) was added and after 10 minutes at room temperature, the cells were added on top of the peptide template at the concentration of 40,000 cells per well. Mitochondrial potential loss was measured using the Varioskan TM kinetic plate reader at excitation 545 nm and emission 590 nm for 3 hours at 27 degrees (kinetic measurements every 5 minutes). Mitochondrial depolarization was normalized to DMSO control (0%) and positive control FCCP (100%) (carbonyl cyanide 4-(triflyoromethoxy) phenyl hydrazone).

### High content fluorescent microscopy

This method was used to assess apoptotic cell death. Cells were seeded into a 384-well glass-bottom black plate (IBL, P384-1.5H-N) and left to adhere overnight. After 72 hr treatments, 0.5 μg/ml PI (Sigma-Aldrich, P4864) and 1.4 μg/ml Hoechst 33342 (Invitrogen, H3570) were added to cells. Images were then taken on the ArrayScan XTI Live High Content Platform (ThermoFisher). Drugs were normalised to paired vehicle controls, followed by 0 µM 5FU. GraphPad Prism was used to calculate the area under the curve (AUC) for both parameters ± 5FU and then the difference in AUC was calculated as: (*AUC* + 5*FU*) − (*AUC* − 5*FU*). A negative value for the valid object count signifies synergism of the CI inhibitors and 5FU, with the opposite true for % PI. Growth arrest was determined as 1-average FOC.

### RNA-sequencing

Total RNA was extracted from HCT116 cells and *AKP* organoids using the Roche KAPA RNA HyperPrep kit with RiboErase according to the manufacturer’s protocol. It is designed for NGS library construction from 25 ng–1 μg of total RNA and depletes both cytoplasmic (5S, 5.8S, 18S and 28S) and mitochondrial (12S and16S) ribosomal RNA species. Library QC was performed on the fragment analyser (HS NGS kit) and Qubit. Finally, the libraries were sequenced using the Illumina NovaSeq 6000 sequencing system. The indexes used for these libraries were from the KAPA Unique Dual-Indexed Adapter kit.

### RNA-sequencing data analysis

RNA-seq reads were aligned to the human genome (hg37) using STAR aligner^87^ and the number of reads mapping to genes annotated in Gencode build 37 (v22) were calculated using HTseq^88^. Differentially expressed genes were identified using the DESeq2 R package^89^ and results were visualised as a volcano plot using the ggplot2 R package^90^ (https://ggplot2.tidyverse.org.) Metabolic genes were defined based on a curated list^10,55^. Gene set enrichment analysis was performed using the clusterProfiler^91^ and msigdbr R packages^92^ (https://CRAN.R-project.org/package=msigdbr>). Heatmaps were generated using Morpheus Broad Institute website. All data generated from >2 psuedo-biological replicates per timepoint.

### Real-time PCR

Total RNA was isolated from the cells using the Roche High Pure RNA isolation kit (Roche Life Science, 11828665001) and reverse transcription was performed using the iScript cDNA synthesis kit (Biorad, 1708891) according to the manufacturer’s instructions. Quantitative PCR assays were performed using 480 SYBR Green Master reagent mix (Roche Life Science, 4707516001, murine probes) or TaqMan assays (human ETC genes) on a real-time fluorescence qPCR instrument (LightCycler 480II, Roche Life Science). The expression of relative mRNA was normalized to that of the housekeeping gene 18S. The following primers were used:

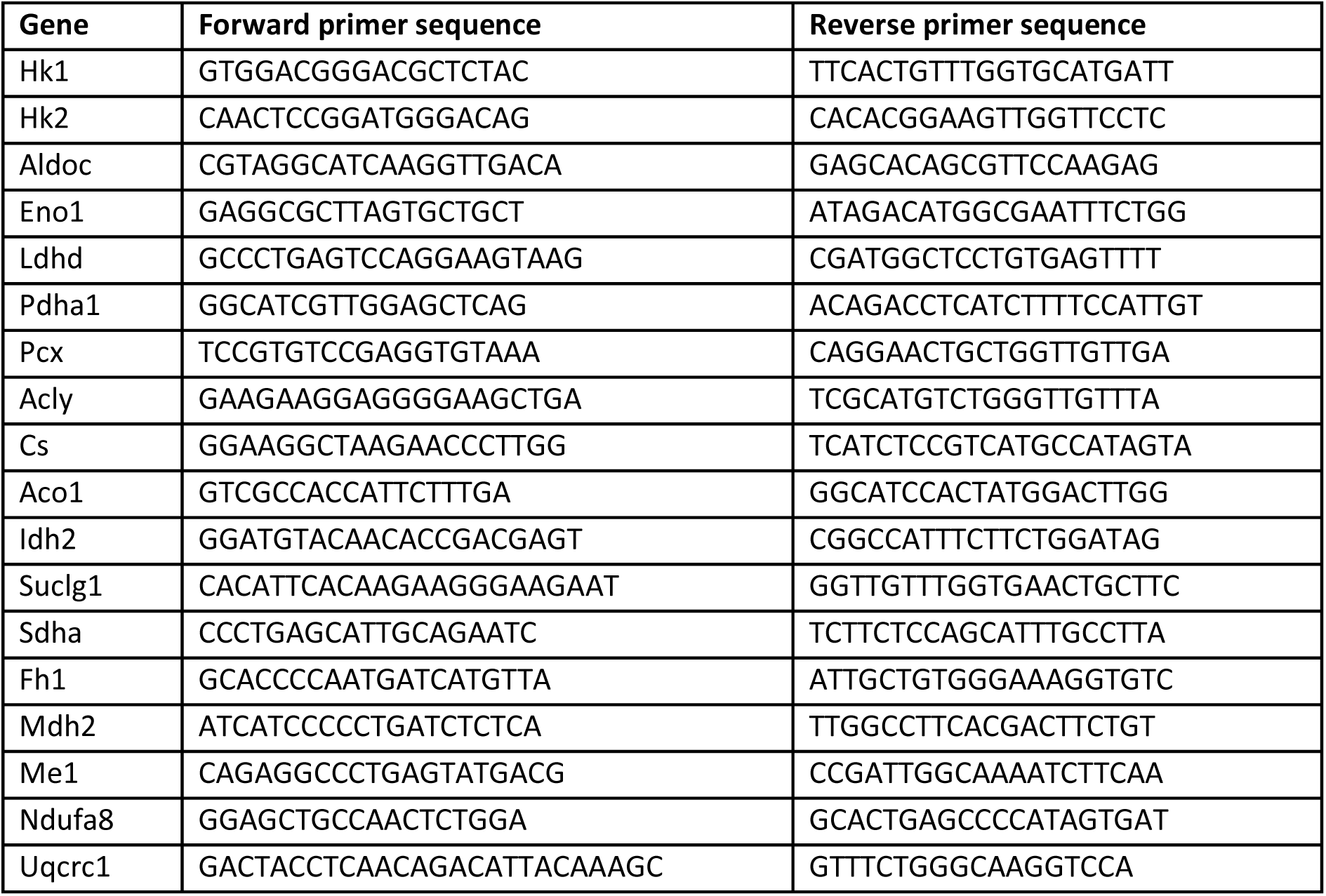

The following TaqMan assays were used:

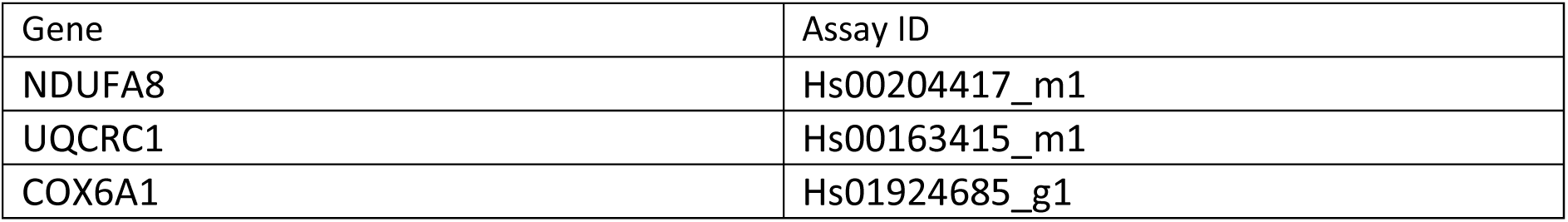

### In vivo studies

Animals were maintained under specific-pathogen-free (SPF) conditions and in compliance with the NI Department of Health (under PPL #2874) regulations with approval from Queen’s University Belfast (QUB) Animal Welfare and Ethical Review Body (AWERB). Mice were housed with no more than 5 animals/cage and were kept on an ad libitum diet. Sample sizes are shown in the figures or legends. Xenograft studies were completed in female mice, all GEMM studies used animals of both sexes in each treatment group.

For human xenograft studies, 2×10^6^ HCT116 cells were transplanted 1:1 in Growth Factor Reduced Matrigel (Corning, CLS356231) into athymic nude mice and volume calculated using formula ½(a x b^2^). When tumours reached ∼100 mm^3^ average per genotype, animals were randomised and given 5FU (10 mg/kg), Metformin (250 mg/kg) or vehicle (PBS) daily via i.p. (5FU) and o.g. (Metformin) for 11 days.

For GEMM studies, intestinal tumours were generated through intraperitoneal (i.p.) injection of 80 mg/kg tamoxifen in 8-10 week old VillinCre^ERT2^;Apc^Fx/+^;Kras^G12D/+^;p53^Fx/Fx^ mice (*A*(Het)*KP* mice or intracolonic induction of colon tumours by submucosal injection of 100 μM 4-OHT (Sigma) in 8-10 week old VillinCre^ERT2^;Apc^Fx/Fx^;Kras^G12D/+^;p53^Fx/Fx^ mice (*AKP*) mice. Treatment was initiated in mice when they displayed clinical signs characteristic of intestinal tumour burden as defined in the relevant licensing documents (e.g. hunching, bloody stools). Mice were randomly assigned to cohorts and received a single dose of indicated compounds: vehicle (PBS) by i.p. and/or oral gavage (o.g.), 250 mg/kg metformin (MET; o.g.) alone, 150 mg/kg 5FU (i.p.) alone or in combination with MET. After 72 hr, mice were culled, and tissue was collected. For survival analysis: *AKP* tumour bearing mice were generated as above, and 26 days after induction they were treated with 150 mg/kg 5FU (i.p.) ± 250 mg/kg Metformin (o.g.) thrice weekly. Animals were culled at clinical endpoint, and survival analysed by Kaplan Meier methods.

### Tissue preparation and staining

Intestinal tissues were flushed with PBS and fixed as a Swiss roll in 10% buffered formalin overnight. These were then transferred to 70% ethanol prior to processing for embedding. All haematoxylin and eosin (H&E) and immunohistochemistry (IHC) staining was performed on 5 μm formalin-fixed paraffin-embedded sections (FFPE) that had previously been heated at 55 °C for 45 minutes.

Standard protocols were used for H&E staining. Slides were deparaffinised and dehydrated, and antigen retrieval was performed using either Tris-EDTA or sodium citrate buffer as recommended by antibody supplier. The slides were then stained with standard tertiary method: incubated with indicated primary antibodies (TOM20 (Abcam, ab186734; 1:200) and Cleaved Caspase 3 (CC3; Cell Signalling Technology, 9661L; 1:200)) at 4 °C overnight; followed by Goat anti-rabbit IgG (H+L) (Vector Laboratories, BA-1000; 1:200) for 1 hr, followed by streptavidin-HRP tertiary antibody (Vector Laboratories, SA-5004; 1:200) for 30 minutes. Samples were further stained with a DAB kit (Abcam, ab64238) and counterstained with haematoxylin. Images were acquired using brightfield microscopy at x20 objective lens and analysed using either QuPath or Image J. We quantified the DAB signal area in the IHC image using Image J as previously described^93^ and calculated the optical density: log (max intensity/min intensity).

### CRISPR

GeCKO Library V1 sgRNA library/pool targeting 17,419 genes was a kind gift from the Zhang laboratory. Library was amplified, lentivirus produced and titred as per Shalem et al.^94^ with adaptations as described previously^95^. Briefly, HCT116 cells were infected with GeCKO v1 human CRISPR knockout pooled library using lentiviral transduction at an MOI of 0.3, selected and populations stably expressing gRNAs were selected for using puromycin for 72 hours and expanded for 8 days to allow for essential gene drop-out. 30×10^6^ cells were harvested (representing an estimated coverage of 300x) and 30×10^6^ cells plated for treatment the next day with 5 μM 5FU, or media only for a further 11 days. Cells were passaged when reached 70-80% confluence again maintaining 30×10^6^ cells at each serial passage, 5FU added in fresh media every 3 days. DNA was extracted and sgRNA inserts amplified as per Lees et al.^95^. Adapters were trimmed from de-multiplexed FASTQ files using Cutadapt (https://pypi.python.org/pypi/cutadapt), aligned to the Gecko V1 design file and count tables generated with MAGeCK software V5. Positively and negatively selected sgRNAs were identified at sgRNA, gene and pathway level using MAGeCK RRA algorithm^96^.

### Taxonomy cohort

The gene expression series matrix was downloaded directly from the Gene Expression Omnibus (GEO) using accession number GSE103479. The datasets had already been Robust Multi-array Average (RMA)-normalised in batches using makecdfenv, affy and limma R packages, and batch corrected using batch the ComBat method in the sva R package.

Patients were subdivided by patients who received surgery alone or adjuvant 5FU-based chemotherapy and subsequently grouped based on the median overall survival (OS) into either ‘good outcome’ or ‘poor outcome’. OS comparisons were conducted by Kaplan-Meier analysis using Log-rank testing for patients separated based on the mean OxPhos score generated using ssGSEA.

### Statistical Analysis

Data was visualised and statistical analyses performed using Prism 9.5.1 software (Graph Pad). P<0.05 was considered statistically significant. In all cases, experimental groups showed comparable variance. P values for unpaired comparisons between two groups with comparable variance were calculated by two-tailed Student’s t-test. One-way ANOVA with Dunnett’s or Tukey’s multiple comparisons was used for analysis between datasets with one independent variable. Ordinary two-way ANOVA was used for analysis that involved two independent variables, followed by Sidak’s or Tukey’s post-test for individual comparisons. Kaplan Meier comparison was used for analysis of survival cohorts. RNA-Seq gene expression data was analysed using negative binomial generalised linear model (DESeq2). *P<0.05, **P<0.01, ***P<0.001, ****P<0.0001. Error bars indicate mean ±s.d. or s.e.m., as indicated.

## Data availability

All RNA-seq data discussed in this publication have been deposited in NCBI’s Gene Expression Omnibus^97^ and are accessible through GEO Series accession numbers GSE272456 (HCT116) and GSE272391 (AKP organoids).

## Contributions

**Conceived and designed the analysis**: DYM, CMC, TNC, SSMD, DBL, IGM, MJLB, EMK; **Collected the data**: DYM, CMC, NL, CB, AP, SG, ER, WJMD, ARoe, AS, CO, FF, MD, AJE, LMS, PFG, AB, SP, BC, AYC; **Contributed data or analysis tools:** SBM, PDD, SP, BG, CE, ARyan, SSMD, OS, TNC, DBL; **Performed the analysis:** DYM, CMC, NL, CB, AP, SG, ER, WJMD, ARoe, AYC, AS, CO, RMC, SM, MD, AJE, LMS, SSMD, EMK; **Wrote the paper:** DYM, DBL, SSMD, MJLB, EMK; **Other**: All authors contributed to editing process.

## Acknowledgements

We thank the patients who contributed to the datasets used in the analyses presented. We thank the Biological Services Unit at Queen’s University Belfast for supporting the animal studies undertaken. This work was supported by Cancer Research UK (C61288/A26045).

## Footnotes

fn Global estimate of patients treated each year with a TS-targeted therapy is based on cancer incidence data obtained from GLOBOCAN 2020, the Global Cancer Observatory Cancer Incidence and Mortality Worldwide, the World Health Organization, SEER, www.cancer.gov (National Institutes of Health) and www.nhs.gov.uk (National Health Service) and adjusted by clinical parameters and evidence-based treatment guidelines to estimate the number of newly diagnosed cancer cases that have a high probability of being treated with a TS-targeted therapy.

## References

1 Hlavata, I. et al. The role of ABC transporters in progression and clinical outcome of colorectal cancer. Mutagenesis 27, 187–196 (2012).

2 Zhang, L. et al. Tumor-associated macrophages confer colorectal cancer 5-fluorouracil resistance by promoting MRP1 membrane translocation via an intercellular CXCL17/CXCL22–CCR4– ATF6–GRP78 axis. Cell Death & Disease 14, 582 (2023).

3 Khan, S. U., Fatima, K., Aisha, S. & Malik, F. Unveiling the mechanisms and challenges of cancer drug resistance. Cell Communication and Signaling 22, 109 (2024).

4 Haller, D. G. et al. Capecitabine plus oxaliplatin compared with fluorouracil and folinic acid as adjuvant therapy for stage III colon cancer. Journal of clinical oncology 29, 1465–1471 (2011).

5 Kuebler, J. P. et al. Oxaliplatin combined with weekly bolus fluorouracil and leucovorin as surgical adjuvant chemotherapy for stage II and III colon cancer: results from NSABP C-07. Journal of clinical oncology 25, 2198–2204 (2007).

6 André, T. et al. Improved overall survival with oxaliplatin, fluorouracil, and leucovorin as adjuvant treatment in stage II or III colon cancer in the MOSAIC trial. Journal of clinical oncology 27, 3109–3116 (2009).

7 Douillard, J. Y. et al. Irinotecan combined with fluorouracil compared with fluorouracil alone as first-line treatment for metastatic colorectal cancer: a multicentre randomised trial. The Lancet 355, 1041–1047 (2000).

8 Longley, D. B., Harkin, D. P. & Johnston, P. G. 5-fluorouracil: mechanisms of action and clinical strategies. Nature reviews cancer 3, 330–338 (2003).

9 Ser, Z. et al. Targeting one carbon metabolism with an antimetabolite disrupts pyrimidine homeostasis and induces nucleotide overflow. Cell reports 15, 2367–2376 (2016).

10 Gaude, E. & Frezza, C. Tissue-specific and convergent metabolic transformation of cancer correlates with metastatic potential and patient survival. Nature communications 7, 13041 (2016).

11 Hanahan, D. Hallmarks of cancer: new dimensions. Cancer discovery 12, 31–46 (2022).

12 Lee, S. et al. Targeting mitochondrial oxidative phosphorylation abrogated irinotecan resistance in NSCLC. Scientific reports 8, 15707 (2018).

13 Chen, C. et al. Oxidative phosphorylation enhances the leukemogenic capacity and resistance to chemotherapy of B cell acute lymphoblastic leukemia. Science advances 7, eabd6280 (2021).

14 Evans, K. W. et al. Oxidative phosphorylation is a metabolic vulnerability in chemotherapy- resistant triple-negative breast cancer. Cancer research 81, 5572–5581 (2021).

15 Ludikhuize, M. C. et al. Rewiring glucose metabolism improves 5-FU efficacy in p53- deficient/KRAS^G12D^ glycolytic colorectal tumors. Communications Biology 5, 1159 (2022).

16 Liang, Y. et al. Dichloroacetate restores colorectal cancer chemosensitivity through the p53/miR-149-3p/PDK2-mediated glucose metabolic pathway. Oncogene 39, 469–485 (2020).

17 Tadros, S. et al. De novo lipid synthesis facilitates gemcitabine resistance through endoplasmic reticulum stress in pancreatic cancer. Cancer research 77, 5503–5517 (2017).

18 Rashmi, R. et al. Radioresistant cervical cancers are sensitive to inhibition of glycolysis and redox metabolism. Cancer research 78, 1392–1403 (2018).

19 Birsoy, K. et al. An essential role of the mitochondrial electron transport chain in cell proliferation is to enable aspartate synthesis. Cell 162, 540–551 (2015).

20 Bock, F. J. & Tait, S. W. G. Mitochondria as multifaceted regulators of cell death. Nature reviews molecular cell biology 21, 85–100 (2020).

21 Vakifahmetoglu-Norberg, H., Ouchida, A. T. & Norberg, E. The role of mitochondria in metabolism and cell death. Biochemical and biophysical research communications 482, 426–431 (2017).

22 Gomes, L. C., Di Benedetto, G. & Scorrano, L. During autophagy mitochondria elongate, are spared from degradation and sustain cell viability. Nature cell biology 13, 589–598 (2011).

23 Crowell, P. D. et al. MYC is a regulator of androgen receptor inhibition-induced metabolic requirements in prostate cancer. Cell Reports 42, 113221 (2023).

24 Mishra, P. & Chan, D. C. Metabolic regulation of mitochondrial dynamics. Journal of Cell Biology 212, 379–387 (2016).

25 Fraser, C., Ryan, J. & Sarosiek, K. BH3 profiling: a functional assay to measure apoptotic priming and dependencies. Methods molecular biology 1877, 61–76 (2019).

26 Dowling, C. M. et al. Multiple screening approaches reveal HDAC6 as a novel regulator of glycolytic metabolism in triple-negative breast cancer. Science advances 7, eabc4897 (2021).

27 Ahmed, D. et al. Epigenetic and genetic features of 24 colon cancer cell lines. Oncogenesis 2, e71 (2013).

28 Eshleman, J. R. et al. Chromosome number and structure both are markedly stable in RER colorectal cancers and are not destabilized by mutation of p53. Oncogene 17, 719–725 (1998).

29 Moss, D. Y., McCann, C. & Kerr, E. M. Rerouting the drug response: Overcoming metabolic adaptation in KRAS-mutant cancers. Science Signaling 15, eabj3490 (2022).

30 Kerr, E. M., Gaude, E., Turrell, F. K., Frezza, C. & Martins, C. P. Mutant Kras copy number defines metabolic reprogramming and therapeutic susceptibilities. Nature 531, 110–113 (2016).

31 Najumudeen, A. K. et al. The amino acid transporter SLC7A5 is required for efficient growth of KRAS-mutant colorectal cancer. Nature genetics 53, 16–26 (2021).

32 Son, J. et al. Glutamine supports pancreatic cancer growth through a KRAS-regulated metabolic pathway. Nature 496, 101–105 (2013).

33 Vande Voorde, J., et al. Metabolic profiling stratifies colorectal cancer and reveals adenosylhomocysteinase as a therapeutic target. Nature metabolism 5, 1303–1318 (2023).

34 Jiang, P. et al. p53 regulates biosynthesis through direct inactivation of glucose-6-phosphate dehydrogenase. Nature cell biology 13, 310–316 (2011).

35 Vernucci, E. et al. Metabolic alterations in pancreatic cancer progression. Cancers 12, 2 (2019).

36 Oni, T. E. et al. SOAT1 promotes mevalonate pathway dependency in pancreatic cancer. Journal of experimental medicine 217, e20192389 (2020).

37 Sinthupibulyakit, C., Ittarat, W., St Clair, W. H. & St Clair, D. K. p53 Protects lung cancer cells against metabolic stress. International journal of oncology 37, 1575–1581 (2010).

38 Jackstadt, R. et al. Epithelial NOTCH signaling rewires the tumor microenvironment of colorectal cancer to drive poor-prognosis subtypes and metastasis. Cancer cell 36, 319–336. e317 (2019).

39 Jackstadt, R. & Sansom, O. J. Mouse models of intestinal cancer. The Journal of pathology 238, 141–151 (2016).

40 Tidwell, T. R., Røsland, G. V., Tronstad, K. J., Søreide, K. & Hagland, H. R. Metabolic flux analysis of 3D spheroids reveals significant differences in glucose metabolism from matched 2D cultures of colorectal cancer and pancreatic ductal adenocarcinoma cell lines. Cancer & Metabolism 10, 9 (2022). 10.1186/s40170-022-00285-w

41 Davidson, S. M. et al. Environment impacts the metabolic dependencies of Ras-driven non-small cell lung cancer. Cell metabolism 23, 517–528 (2016).

42 Roper, J. et al. In vivo genome editing and organoid transplantation models of colorectal cancer and metastasis. Nature biotechnology 35, 569–576 (2017).

43 Roper, J. et al. Colonoscopy-based colorectal cancer modeling in mice with CRISPR–Cas9 genome editing and organoid transplantation. Nature protocols 13, 217–234 (2018).

44 Vaquero-Siguero, N. et al. Modeling Colorectal Cancer Progression Reveals Niche-Dependent Clonal Selection. Cancers 14, 4260 (2022).

45 Jackman, A. L. et al. ICI D1694, a quinazoline antifolate thymidylate synthase inhibitor that is a potent inhibitor of L1210 tumor cell growth in vitro and in vivo: a new agent for clinical study. Cancer Research 51, 5579–5586 (1991).

46 Argilés, G. et al. Localised colon cancer: ESMO Clinical Practice Guidelines for diagnosis, treatment and follow-up. Annals of oncology 31, 1291–1305 (2020).

47 NICE. Colorectal cancer, <https://www.nice.org.uk/guidance/ng151> (2020).

48 Blondy, S. et al. 5-Fluorouracil resistance mechanisms in colorectal cancer: From classical pathways to promising processes. Cancer science 111, 3142–3154 (2020).

49 Hammond, W. A., Swaika, A. & Mody, K. Pharmacologic resistance in colorectal cancer: a review. Therapeutic advances in medical oncology 8, 57–84 (2016).

50 Rossignol, R. et al. Energy substrate modulates mitochondrial structure and oxidative capacity in cancer cells. Cancer research 64, 985–993 (2004).

51 Orlicka-Płocka, M., Gurda-Wozna, D., Fedoruk-Wyszomirska, A. & Wyszko, E. Circumventing the Crabtree effect: Forcing oxidative phosphorylation (OXPHOS) via galactose medium increases sensitivity of HepG2 cells to the purine derivative kinetin riboside. Apoptosis 25, 835–852 (2020).

52 Boyer, J. et al. Characterization of p53 wild-type and null isogenic colorectal cancer cell lines resistant to 5-fluorouracil, oxaliplatin, and irinotecan. Clinical cancer research 10, 2158–2167 (2004).

53 Allen, W. L. et al. Transcriptional subtyping and CD8 immunohistochemistry identifies poor prognosis stage II/III colorectal cancer patients who benefit from adjuvant chemotherapy. JCO precision oncology 2, 1–15 (2018).

54 Liberzon, A. et al. Molecular signatures database (MSigDB) 3.0. Bioinformatics 27, 1739–1740 (2011).

55 Liberzon, A. et al. The molecular signatures database hallmark gene set collection. Cell systems 1, 417–425 (2015).

56 Molina, J. R. et al. An inhibitor of oxidative phosphorylation exploits cancer vulnerability. Nature medicine 24, 1036–1046 (2018).

57 Yap, T. A. et al. Complex I inhibitor of oxidative phosphorylation in advanced solid tumors and acute myeloid leukemia: phase I trials. Nature medicine 29, 115–126 (2023).

58 NICE. Type 2 diabetes in adults: management, <https://www.nice.org.uk/guidance/ng28/chapter/Recommendations#first-line-drug-treatment> (2015).

59 Owen, M. R., Doran, E. & Halestrap, A. P. Evidence that metformin exerts its anti-diabetic effects through inhibition of complex 1 of the mitochondrial respiratory chain. Biochemical journal 348, 607–614 (2000).

60 Hirsch, A. et al. Metformin inhibits human androgen production by regulating steroidogenic enzymes HSD3B2 and CYP17A1 and complex I activity of the respiratory chain. Endocrinology 153, 4354–4366 (2012).

61 Bridges, H. R., Jones, A. J. Y., Pollak, M. N. & Hirst, J. Effects of metformin and other biguanides on oxidative phosphorylation in mitochondria. Biochemical journal 462, 475–487 (2014).

62 Bankhead, P. et al. QuPath: Open source software for digital pathology image analysis. Scientific reports 7, 16878

63 Vasan, N., Baselga, J. & Hyman, D. M. A view on drug resistance in cancer. Nature 575, 299–309 (2019). 10.1038/s41586-019-1730-1

64 Masoud, R. et al. Targeting mitochondrial complex I overcomes chemoresistance in high OXPHOS pancreatic cancer. Cell reports medicine 1, 100143 (2020).

65 Farge, T. et al. Chemotherapy-resistant human acute myeloid leukemia cells are not enriched for leukemic stem cells but require oxidative metabolism. Cancer discovery 7, 716–735 (2017).

66 Wong, C. C. et al. In colorectal cancer cells with mutant KRAS, SLC25A22-mediated glutaminolysis reduces DNA demethylation to increase WNT signaling, stemness, and drug resistance. Gastroenterology 159, 2163–2180. e2166 (2020).

67 Källberg, J. et al. Intratumor heterogeneity and cell secretome promote chemotherapy resistance and progression of colorectal cancer. Cell Death & Disease 14, 306 (2023).

68 Pranzini, E. et al. SHMT2-mediated mitochondrial serine metabolism drives 5-FU resistance by fueling nucleotide biosynthesis. Cell reports 40, 111233 (2022).

69 Montrose, D. C. et al. Exogenous and endogenous sources of serine contribute to colon cancer metabolism, growth, and resistance to 5-fluorouracil. Cancer research 81, 2275–2288 (2021).

70 Chen, J. et al. The loss of SHMT2 mediates 5-fluorouracil chemoresistance in colorectal cancer by upregulating autophagy. Oncogene 40, 3974–3988 (2021).

71 Huang, C. Y. et al. Glucose metabolites exert opposing roles in tumor chemoresistance. Frontiers in oncology 9, 1282 (2019).

72 Wang, T. et al. Glycolysis is essential for chemoresistance induced by transient receptor potential channel C5 in colorectal cancer. BMC cancer 18, 207 (2018).

73 Sainero-Alcolado, L., Liaño-Pons, J., Ruiz-Pérez, M. V. & Arsenian-Henriksson, M. Targeting mitochondrial metabolism for precision medicine in cancer. Cell Death & Differentiation 29, 1304–1317 (2022).

74 Kwong, S. C. & Brubacher, J. Phenformin and lactic acidosis: a case report and review. The Journal of emergency medicine 16, 881–886 (1998).

75 Vernieri, C. et al. Impact of Metformin Use and Diabetic Status During Adjuvant Fluoropyrimidine-Oxaliplatin Chemotherapy on the Outcome of Patients with Resected Colon Cancer: A TOSCA Study Subanalysis. Oncologist 24, 385–393 (2019).

76 Tarhini, Z. et al. The effect of metformin on the survival of colorectal cancer patients with type 2 diabetes mellitus. Scientific Reports 12, 12374 (2022).

77 Ng, C. A. W. et al. Metformin and colorectal cancer: a systematic review, meta-analysis and meta-regression. International journal of colorectal disease 35, 1501–1512 (2020).

78 Piskounova, E. et al. Oxidative stress inhibits distant metastasis by human melanoma cells. Nature 527, 186–191 (2015).

79 Bezwada, D. et al. Mitochondrial metabolism in primary and metastatic human kidney cancers. *Preprint at* https://www.biorxiv.org/content/10.1101/2023.02.06.527285v1 (2023).

80 Porporato, P. E. et al. A mitochondrial switch promotes tumor metastasis. Cell reports 8, 754–766 (2014).

81 Sun, X. et al. Increased mtDNA copy number promotes cancer progression by enhancing mitochondrial oxidative phosphorylation in microsatellite-stable colorectal cancer. Signal transduction and targeted therapy 3, 8 (2018).

82 National Cancer Institute, D., Surveillance Research Program. (2021).

83 Sung, H., et al. Global cancer statistics 2020: GLOBOCAN estimates of incidence and mortality worldwide for 36 cancers in 185 countries. CA: a cancer journal for clinicians 71, 209–249 (2021).

84 Vichai, V. & Kirtikara, K. Sulforhodamine B colorimetric assay for cytotoxicity screening. Nature protocols 1, 1112–1116 (2006).

85 Foight, G. W., Ryan, J. A., Gullá, S. V., Letai, A. & Keating, A. E. Designed BH3 Peptides with High Affinity and Specificity for Targeting Mcl-1 in Cells. ACS Chemical Biology 9, 1962–1968 (2014). 10.1021/cb500340w

86 Ryan, J. & Letai, A. BH3 profiling in whole cells by fluorimeter or FACS. Methods 61, 156–164 (2013). 10.1016/j.ymeth.2013.04.006

87 Dobin, A. et al. STAR: ultrafast universal RNA-seq aligner. Bioinformatics 29, 15–21 (2013). 10.1093/bioinformatics/bts635

88 Anders, S., Pyl, P. T. & Huber, W. HTSeq--a Python framework to work with high-throughput sequencing data. Bioinformatics 31, 166–169 (2015). 10.1093/bioinformatics/btu638

89 Love, M. I., Huber, W. & Anders, S. Moderated estimation of fold change and dispersion for RNA-seq data with DESeq2. Genome Biol 15, 550 (2014). 10.1186/s13059-014-0550-8

90 Wickham, H. ggplot2: Elegant graphics for data analysis. (Springer-Verlag, 2016).

91 Wu, T. et al. clusterProfiler 4.0: A universal enrichment tool for interpreting omics data. Innovation (Camb) 2, 100141 (2021). 10.1016/j.xinn.2021.100141

92 Dolgalev, I. msigdbr: MSigDB Gene Sets for Multiple Organisms in a Tidy Data Format., <https://CRAN.R-project.org/package=msigdbr>> (2022).

93 Crowe, A. R. & Yue, W. Semi-quantitative determination of protein expression using immunohistochemistry staining and analysis: an integrated protocol. Bio-protocol 9, e3465 (2019).

94 Shalem, O. et al. Genome-scale CRISPR-Cas9 knockout screening in human cells. Science 343, 84–87 (2014). 10.1126/science.1247005

95 Lees, A. et al. The pseudo-caspase FLIP (L) regulates cell fate following p53 activation. Proceedings of the National Academy of Sciences 117, 17808–17819 (2020).

96 Li, W. et al. MAGeCK enables robust identification of essential genes from genome-scale CRISPR/Cas9 knockout screens. Genome biology 15, 1–12 (2014).

97 Edgar, R., Domrachev, M. & Lash, A. E. Gene Expression Omnibus: NCBI gene expression and hybridization array data repository. Nucleic Acids Res 30, 207–210 (2002). 10.1093/nar/30.1.207

